# Intestinal enteroendocrine cell subtype differentiation and hormone production in zebrafish

**DOI:** 10.1101/2025.01.17.633579

**Authors:** Margaret Morash, Richard G. Kay, Erik J. Soderblom, Grace H. MacLean, Jia Wen, Peyton J. Moore, Colin R. Lickwar, Fiona M. Gribble, Frank Reimann, Rodger A. Liddle, John F. Rawls

## Abstract

Enteroendocrine cells (EECs) are rare sensory cells in the intestinal epithelium that coordinate digestive physiology by secreting a diverse repertoire of peptide hormones. These hormones are the main effectors of EEC function, and their characterization requires direct observation by mass spectrometry due to the specialized protein cleavage and posttranslational modifications that yield their mature forms. Based on the distinct subset of hormones they predominantly secrete, EECs can be categorized into subtypes. How each EEC subtype is specified, however, remains poorly understood. Here we describe EEC subtype differentiation and hormone production in the zebrafish. Using single-cell RNA sequencing data, we identified EEC progenitors and six EEC subtypes in zebrafish and revealed that their expression profiles are consistent across larval and adult stages. Mass spectrometry analysis of isolated zebrafish EECs identified highly processed peptides derived from 18 of 21 hormone coding genes expressed by EECs, yielding a catalog of >400 unique EEC hormone peptides. We assembled reporters for zebrafish EEC subtypes to test the lineage relationships between EEC subtypes and the EEC progenitor population, which expresses *neurogenin3*. Despite its essential role in mammalian EEC differentiation, we found that selective cytotoxic ablation of *neurogenin3*+ cells in zebrafish only reduced a subset of EEC subtypes. Finally, we discovered that selective ablation of *ghrelin*+ EECs reduced a different subset of EEC subtypes, together suggesting that *neurogenin3*+ and *ghrelin*+ cells serve as distinct precursors for separate EEC subtypes. We anticipate these observations and resources will facilitate future studies in the zebrafish to discern the developmental biology, physiology, and endocrinology of EEC subtypes.

## Introduction

The ability of the intestine to coordinate digestive physiology in response to ingested food, resident microorganisms, and other stimuli is made possible by contributions from specialized sensory epithelial cells called enteroendocrine cells (EECs). EECs are an ancient and conserved feature of the digestive tract in bilaterian animals [1–3], with evidence of cells with similar function in distantly related placozoa [4,5]. EECs respond to stimuli by releasing secretory vesicles containing at least one of over a dozen different hormones. These EEC hormones govern diverse physiologic functions like nutrient digestion, gut motility, satiety, and insulin signaling [6,7]. Each EEC peptide hormone is produced through the transcription and translation of its preprohormone protein, which is then subjected to successive cleavage and posttranslational modification steps prior to secretion from the cell [8]. These highly specialized processing steps make it largely impossible to predict the final functional peptide’s form from the primary amino acid sequence alone, necessitating direct observation of peptides through sensitive peptidomic studies to shed light on their biology.

As EEC peptide hormone functions are highly varied and sometimes antagonistic, ensuring the release of the appropriate hormone is essential for mounting the appropriate physiologic response to a given stimulus. This is largely accomplished through the differentiation of EECs into distinct subtypes that each secrete a single or small subset of hormones [6,7]. EEC differentiation begins with secretory progenitor cells within the intestinal epithelium, which can give rise to any secretory lineage, such as enteroendocrine, goblet, tuft, and Paneth cells [9]. In mouse and humans, transient expression of the transcription factor *Neurogenin3* (*Ngn3*) commits these secretory progenitors to an EEC fate, and *Ngn3* expression is therefore used as a marker of EEC progenitors [10–14]. Subsequent expression of select transcription factors including *Neuronal differentiation 1* (*Neurod1)* are important in mammals for the differentiation of *Ngn3*+ EEC progenitors into mature EEC subtypes and are expressed broadly across EEC subtypes [15–18]. More recent studies in mammalian organoid models have proposed successive factors expressed only in a subset of EECs that drive EECs towards a single specific subtype fate [19,20]. This work has begun to shed light on how EEC subtype identity – and thus the cell’s hormone profile and function – is determined, but further studies and *in vivo* testing are needed to elucidate this important process.

While EEC subtypes were historically believed to be terminally differentiated and therefore insulated from one another once committed, recent work in organoids indicated that some EEC subtypes can morph into others as the cells travel up the crypt-villus axis [21]. This new concept of EEC subtype plasticity raises interesting questions about the lineage relationships between EEC subtypes. While a new concept in the intestine, there are examples of plastic lineage relationships between endocrine subtypes in other tissues. Studies in the stomach and pancreatic islet show some endocrine subtypes, specifically those producing the hormone ghrelin, can give rise to others [22]. Multiple single cell RNA sequencing (scRNA-seq) studies in mouse have hinted at a similar role for *Ghrelin* (*Ghrl*) expressing cells in the small intestine as they identified a *Ghrl*-expressing EEC progenitor population [23–25]. RNA velocity and partition-based graph abstraction of one of those datasets predicted a differentiation trajectory through the *Ghrl*+ progenitor cluster to many EEC subtypes [24], but a role for *Ghrl*+ cells in EEC differentiation has never been directly tested. Together these studies suggest that establishment of EEC subtype identity may be more dynamic than previously thought. This may be true at the level of an individual EEC, such as a *Ghrelin*-expressing cell possibly giving rise to other EEC subtypes, as well as at the level of the whole organism, such as possible changes to the overall EEC subtype profile as an animal develops. These aspects of EEC subtype dynamics have not been fully investigated and represent an important gap in our understanding of EEC biology.

The zebrafish is an ideal vertebrate model to study EEC subtype dynamics. We previously revealed extensive similarities between EEC morphology, sensitivity, and neural connectivity in zebrafish and mammals [26,27]. Many important aspects of EEC differentiation are conserved between zebrafish, mammals, and insects [28–33], although there are slight differences.

Specifically, *ngn3*, the zebrafish ortholog of *Ngn3*, is expressed in some zebrafish EECs, but is thought to play a smaller role in their differentiation, although its exact function remains unclear [28,30]. Instead, *neurod1*, the zebrafish ortholog of *Neurod1*, has been shown to be necessary and sufficient for EEC differentiation [26,27,31]. Similar to mammals, *neurod1* is expressed across all EECs in zebrafish and is used as a pan-EEC reporter [26]. Critically, the optical transparency of zebrafish enables *in vivo* imaging of gut anatomy and physiology through early adulthood. Combined with the ease of generating genetic reporters, this provides a powerful system for investigating EEC subtype development and dynamics in their native context.

However, zebrafish EEC subtypes and their hormone products have not been previously described.

In this study, we aimed to define EEC subtypes in zebrafish and establish an experimental system in which to study their development and function. To do this, we leveraged scRNA-seq data from larval and adult zebrafish to reveal seven distinct EEC populations, six of which we characterized *in vivo* across larval development. We also deployed peptidomic analysis to demonstrate that these EECs produce a wide array of highly processed and conserved peptide hormones. To better understand how these EEC subtypes develop, we probed their lineage relationships by genetically restricting certain cell states and assessing the impacts on the remaining subtype populations. In this way we were able to reveal that both predicted contributors, such as *ngn3*+ cells, as well as untested factors, such as *ghrl*+ cells, play major roles in development of EEC subtypes in zebrafish.

## Results

### Zebrafish EEC transcriptional signatures separate EEC progenitors and six distinct EEC subtypes

To understand EEC subtype diversity and differentiation in zebrafish, we used established markers of secretory cells [9,28,29,34–47] to identify and subset these populations from published intestinal scRNA-seq datasets derived from larval [48] and adult [49] zebrafish (S1A, S1B Fig). Integration of these secretory cells from larval and adult datasets showed the expected structure of a hub of secretory progenitor cells with spokes of distinct secretory populations (S1C, S1D Fig). With this confirmation in place, we then subsetted and reintegrated just EECs and secretory progenitors into a new combined larval and adult dataset (Fig 1A, 1B). Within this dataset we identified clusters 0-6 as EECs based on *neurod1* expression (Fig 1C) and clusters 7-10 as secretory progenitors (S2A Fig). We found that each EEC cluster contained both larval and adult cells with no apparent age dependency for any EEC subtype (Figs 1D and S2B). We also observed that cells from the larval and adult datasets showed similar distributions of cells across EEC clusters (S2C Fig). These similarities despite the many differences in samples used to generate each dataset (S2G Fig) establish that EEC subtypes and their underlying gene expression programs are surprisingly consistent between larval and mature adult stages in zebrafish.

**Fig 1.**
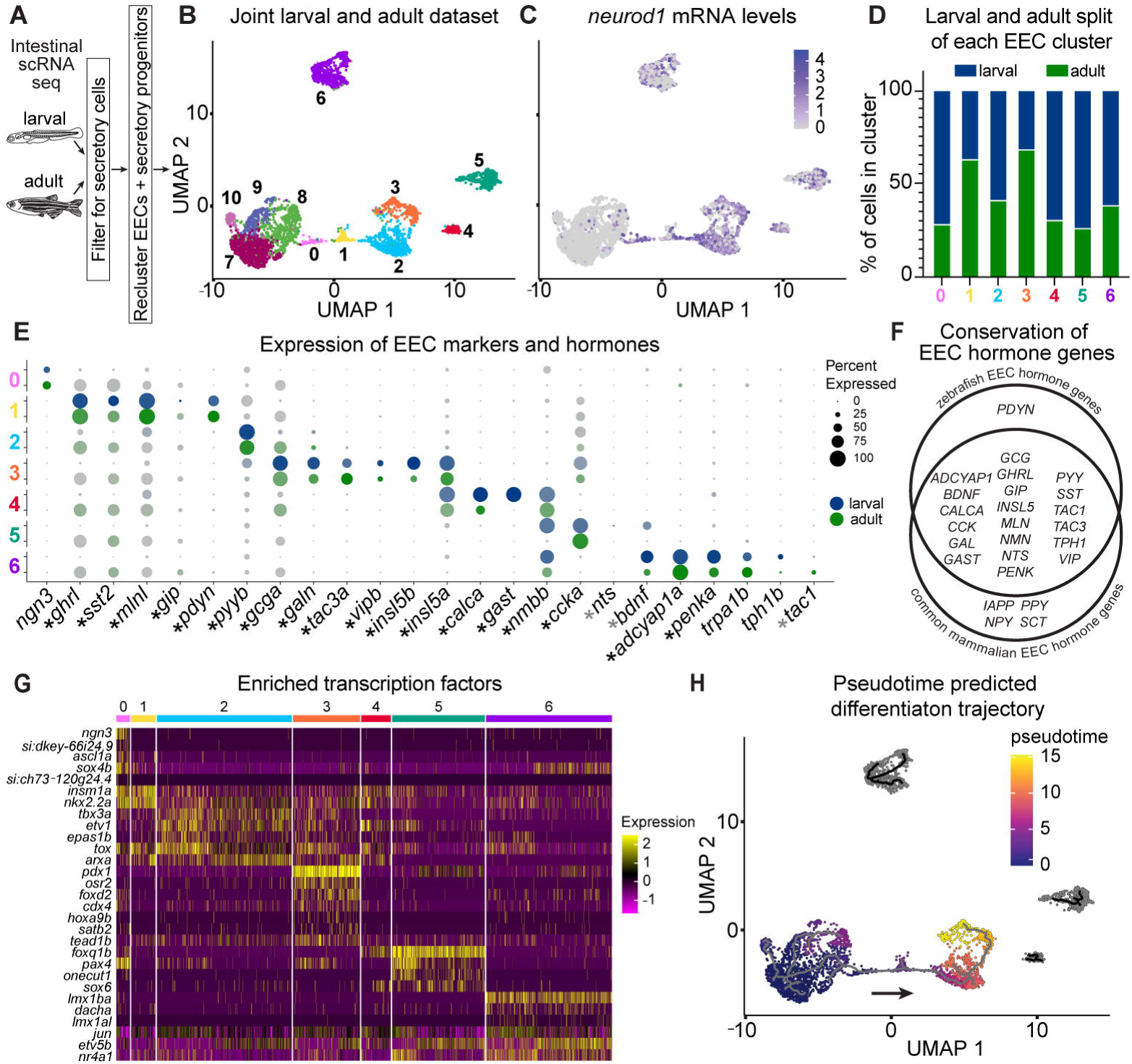
scRNA-seq of larval and adult EECs identifies 7 distinct clusters. **(A)** Schematic overview identifying EECs and secretory cells from published larval [48] and adult [49] datasets. **(B)** UMAP of secretory progenitors and EECs from larval and adult zebrafish identifies 11 clusters. **(C)** Normalized expression of the zebrafish pan-EEC transcription factor and reporter *neurod1* labels clusters 0-6. **(D)** The percentage of cells derived from either the larval (blue) or adult (green) datasets in each of the 7 EEC clusters. **(E)** Expression dotplot of relevant marker genes of the 7 EEC clusters. EEC hormone coding genes are marked with an asterisk next to their gene name on the x axis. Black asterisks indicate genes we identified peptides from and gray asterisks mark those with no detected peptides (Fig 2B and S3). Dots represent gene expression in the specified cluster. Dot radius corresponds to the percent of cells expressing the gene and dot transparency corresponds to expression intensity. Expression is reported for both the larval (blue) and adult (green) cells in the cluster. Note, *gastrin* was not annotated in the adult dataset. **(F)** Venn diagram of EEC hormone coding gene homologs found in zebrafish and mammals. **(G)** Heatmap of expression of transcription factors significantly enriched in each EEC cluster in both larval and adult cells. **(H)** UMAP colored by pseudotime as determined by Monocle 3 when cluster 7 was provided as the origin. Brighter coloring indicates “older” cell age and the black line is the predicted differentiation trajectory.

To identify the shared features of larval and adult EECs, we assessed the expression of EEC markers and hormone coding genes in the combined dataset (Fig 1E). Hormone coding genes (marked with an asterisk in Fig 1E) are often used to separate EECs into subtypes as the hormones these genes encode are the main effectors of known EEC functions. In accord, we found that each EEC hormone coding gene is largely partitioned into a single cluster with remarkable consistency across larval and adult stages. All *neurod1*+ clusters expressed EEC hormone coding genes except for cluster 0, which instead expressed *ngn3*. This pattern is consistent with features of EEC progenitors in mammals, which are marked by *Ngn3* expression and have not yet begun to express hormone coding genes [10–14,50]. As such, we operationally defined cluster 0 as a putative EEC progenitor population and clusters 1-6 as mature EEC subtypes. We observed substantial overlap between the EEC hormone coding genes expressed in our zebrafish dataset and those reported in published mammalian datasets (Fig 1F) [19,23,24,51–55]. While zebrafish and mammalian EECs are not identical, the remarkable consistency in the overall EEC hormone gene repertoire between zebrafish and mammals highlights the critical role of these signaling molecules during vertebrate evolution.

To explore the broader transcriptional programs driving EEC subtype identity, we identified enriched genes specific to each cluster (S1 Table, S2D Fig). Using existing ontologies [56], we identified transcription factors that were significant across larval and adult cells in each cluster (Fig 1G). We also performed pseudotime analysis [57], a computational method for ordering cells along a hypothetical timeline based on transcriptomic similarity that enables the study of cell differentiation dynamics even in samples collected at a single timepoint. Using the secretory progenitor cluster 7 as the start point, this analysis predicted a differentiation trajectory from secretory progenitor clusters to EEC subtype clusters (Fig 1H). These complementary approaches highlighted certain patterns in EEC populations. For example, we note that putative EEC progenitor cluster 0 is enriched in *ascl1a* and *sox4b* in addition to *ngn3*. While mammalian *Ngn3* is the most canonical EEC progenitor marker, studies have also shown that *Ascl1/ascl1a* and *Sox4/sox4b* homologs label endocrine progenitor populations in mammals [19,24,41,58–60] and zebrafish [28,31,61,62]. EEC subtype cluster 1 is enriched for *insm1a* and *nkx2.2a* whose mammalian homologs *Insm1* and *Nkx2.2* are known to play a role in the differentiation of *Ngn3*+ EEC progenitors into mature EEC subtypes [63–66]. Together with the pseudotime data, these findings support a classification of cluster 0 as EEC progenitors and subtype cluster 1 as potentially an early EEC state. Interestingly, subtype cluster 1 also expresses the EEC hormone coding genes *ghrelin* (*ghrl*), *somatostatin* (*sst2*), *motilin-like* (*mlnl*), and *gastric inhibitory peptide* (*gip*), which are canonically thought to mark a mature EEC population [50,67]. EEC subtype clusters 2 and 3 appear to have a close relationship given their adjacent positions in pseudotime and the high expression of subtype cluster 2 enriched transcription factors in subtype cluster 3. EEC subtype cluster 3 is uniquely enriched for the known endocrine transcription factor *pdx1* [68–70] as well as *osr2*, which has been shown in mammals and zebrafish to be more highly expressed in the mid and distal small intestine [23,71], indicating a potential regional distribution of this subtype. Finally, subtype cluster 6, which expresses the rate limiting enzyme in serotonin synthesis *tph1b* (Fig 1E), is enriched for transcription factors *lmx1ba* and *lmx1al* whose orthologs have been shown to be important for the differentiation of serotonergic neurons and EECs, respectively [65,72,73]. Serotonergic EECs, termed enterochromaffin cells, separate from other EECs in mammalian scRNA-seq studies [19,24,25,74]. Likewise, in our dataset we find that subtype cluster 6 most clearly separates from the remaining EEC populations (Fig 1B), suggesting the divergence between enterochromaffin cells and other EECs is an ancestral trait to fishes and mammals. This *in silico* analysis of scRNA-seq data highlights a limited set of transcription factors that might play a role in promoting EEC differentiation from secretory progenitors into discrete EEC subtypes.

### Zebrafish EECs produce fully processed and modified peptide hormones *in vivo*

EEC peptide hormones are derived from the initial translated preprohormone through several steps of specialized cleavage and posttranslational modification (Fig 2A). To understand what processed peptides are produced by zebrafish EECs, we performed mass spectrometry on sorted EECs from larval and adult samples (see Methods). Of the 21 hormone-coding genes expressed by zebrafish EECs (Fig 1E), we identified peptide fragments aligning to 18 of them (Fig 2B). Genes with no detected peptides included *neurotensin* (*nts*) and *tachykinin precursor 1* (*tac1*), which were very lowly expressed at the transcript level, and *brain-derived neurotrophic factor* (*bdnf*), whose peptide product is too large for optimal detection with our methodology. Of note, while expression of *tph1b* is used as a proxy for serotonin because it catalyzes the rate limiting step in its formation, serotonin itself is derived from a single amino acid and is not suited to detection with our methodology. The EEC peptides we identified include products of hormone coding genes specifically expressed by EEC subtypes 1-6, suggesting each of those subtypes produce peptide hormones. We found a modest correlation between the expression level of a gene in scRNA-seq data (S1 Table) and the number of peptides detected that align to that gene (Fig 2C). We also identified peptides from other common components of the hormone-containing secretory vesicles released by EECs. Namely, we detected peptides from *proprotein convertase subtilisin/kexin type 1* and *2* (*pcsk1* and *pcsk2*), known for their role in specialized cleavage of peptide hormones [75], and *secretogranin 2a, 2b, 3,* and *5*, which help form the secretory vesicles and can be processed into hormones themselves (S3Q-S3X Fig) [76–79].

**Fig 2.**
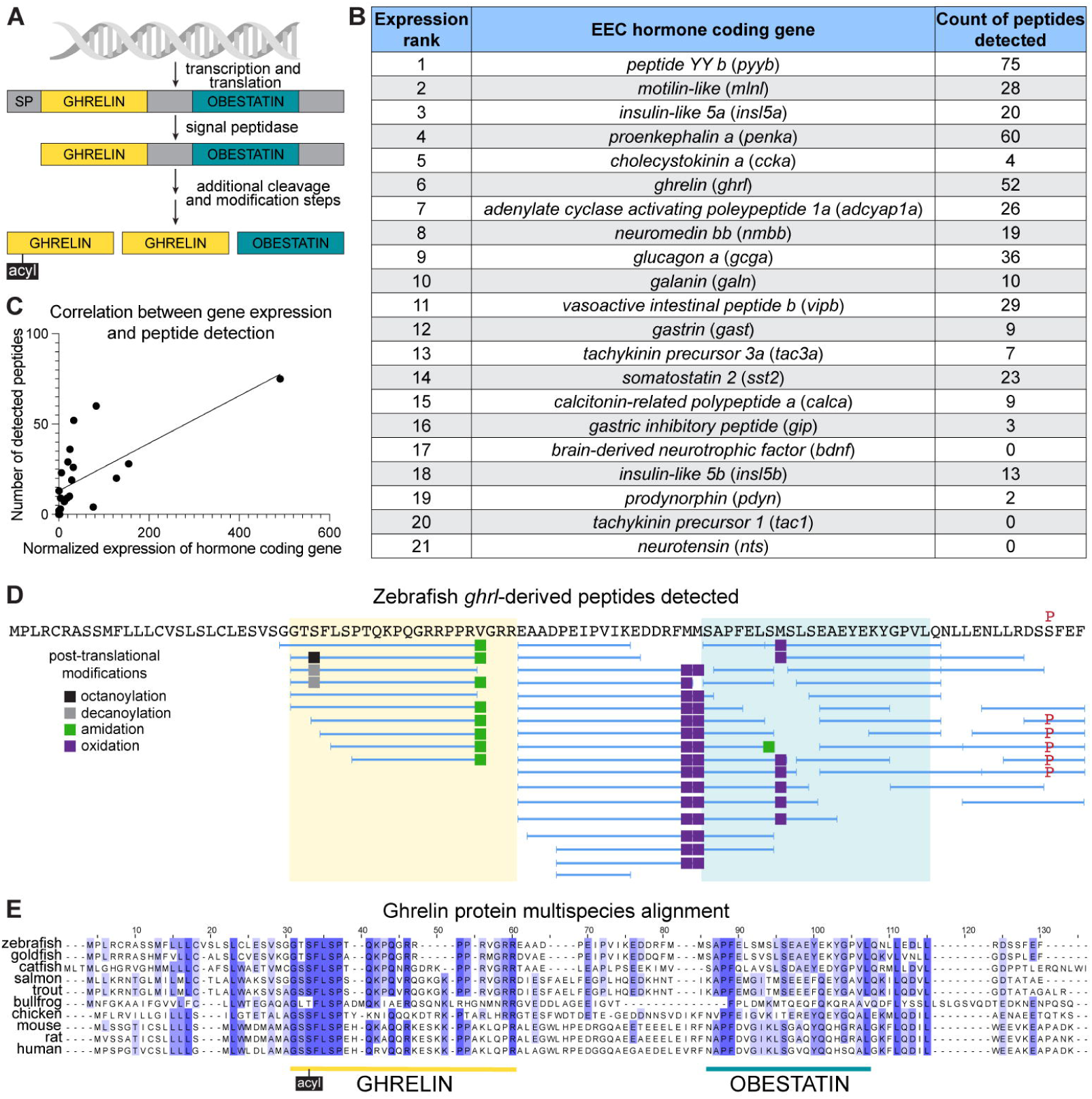
Peptidomic analysis of zebrafish EECs reveals production of conserved and novel peptides. **(A)** Schematic of peptide hormone processing using the human EEC hormone coding gene *GHRELIN* as an example. **(B)** Number of peptides identified from each zebrafish hormone coding gene ranked by normalized mRNA expression level as assessed by scRNA-seq (S1 Table). **(C)** Correlation between the normalized mRNA expression of zebrafish EEC hormone coding genes and the number of peptides detected. **(D)** Representation of *ghrl-*derived peptides detected in zebrafish EECs. The primary amino acid sequence is shown at the top in black with missense variants indicated in red above. Blue horizontal lines below the amino acid sequence represent the individual peptides detected in our study with small vertical lines denoting the stop and start of each peptide. Different colored squares represent various posttranslational modifications detected. Yellow shading marks the region aligning to the functional ghrelin peptide in humans and teal shading marks the region aligning to the functional obestatin peptide in humans. **(E)** Alignment of the primary amino acid sequence of the Ghrelin protein in zebrafish (Uniprot F1QKX9), goldfish (Uniprot Q8AUU1), catfish (Ensembl ENSIPUT00015057246.1), salmon (Uniprot B3IYK1), trout (Uniprot A0A060WHA4), bullfrog (Uniprot Q90W22), chicken (Uniprot Q8AV73), mouse (Uniprot Q9EQX0), rat (Uniprot Q9QYH7), human (Uniprot Q9UBU3). Amino acids are color coded based on their percent identity match across all the reported species with darker coloring indicating a more conserved residue. Human processed peptide annotations are labeled below, including the conserved acylation at the serine 3 residue.

Together, these observations indicate that our EEC peptidomic approach effectively isolated and defined peptide contents of zebrafish EEC secretory vesicles.

Using this dataset, we generated an atlas of zebrafish EEC peptides including their sequence, posttranslational modifications, and alignment to homologous proteins in humans and other vertebrates. This is provided as a resource in the supplemental materials supporting this article (S3 Fig), and we highlight Ghrelin-derived peptides as an example (Fig 2D). In mammals, two main peptide hormone products from the *GHRELIN* gene have been described, ghrelin and obestatin [80,81]. Ghrelin is a 28 amino acid peptide that is often fatty acylated at the serine 3 residue [82]. While both acylated and deacylated ghrelin peptides are produced, fatty acylation improves binding to ghrelin’s receptor and thus is critical to its function [80,82]. As such, the serine 3 residue is highly conserved across species with evidence of acylation at this site in each vertebrate examined (Fig 2 E) [80,83,84]. Indeed, studies in goldfish, a close relative of zebrafish, identified a functional 17 amino acid ghrelin peptide that is acylated at the serine 3 residue [83,85]. Obestatin is a 22 amino acid peptide that was identified more recently by bioinformatic prediction [81]. While initially described as acting in opposition to ghrelin, its function is still debated [81,86]. We detected 35 Ghrelin-derived peptides in zebrafish, including eight that align with the portion of the protein shown to encode a functional ghrelin peptide in humans and goldfish (Fig 2 D, E) [85,87,88]. As fatty acylation is key to ghrelin’s function and not included in the posttranslational modification parameters of our initial search, we performed a second search for just ghrelin peptides and included octanoyl and decanoyl modifications in our parameters. Peptides with both octanoyl-and decanoyl-modified serine 3 residues were detected, demonstrating remarkable conservation of the processing and posttranslational modification of this peptide hormone (Figs 2D, S4A-S4B). While functional studies remain to be performed, the alignment and modification data suggest these peptides could be endogenous zebrafish ghrelin. In contrast, while we detected peptides aligning to portions of the reported obestatin region, we did not detect any peptide spanning that region. Our data, along with the absence of any previous report of an obestatin peptide in other fish species, suggests obestatin may not be present in fishes. We did, however, observe several peptides that align to the unannotated region between the ghrelin and obestatin annotations. There is modest conservation of the amino acids in this region across fishes, but poor conservation between fishes and mammals (Fig 2E), raising the possibility of a peptide produced by EECs in fish but not mammals. Functional studies are needed to test the possible activity of the peptides we detected in this region. Additional examples of conservation and potential novelty in zebrafish EEC peptides are included in the supplemental resource (S3 Fig).

### Zebrafish EEC subtypes show distinct expression patterns *in vivo*

To understand how the seven zebrafish EEC populations we identified develop and function *in vivo*, we assembled reporters for the EEC progenitor population and five of the EEC subtypes (Fig 3A): *ngn3* and its reporter Tg(*ngn3:QF2*) for cluster 0, *ghrl* and its reporter Tg(*ghrl:*QF2) for cluster 1, *peptide YYb* (*pyyb*) and anti-PYY antibody for cluster 2, *glucagon a* (*gcga*) and its reporter Tg(*gcga:GFP*) for cluster 3, *cholecystokinin a* (*ccka*) and anti-CCK antibody for cluster 5, and *transient receptor potential cation channel, subfamily A, member 1b* (*trpa1b*) and its reporter Tg(*trpa1b:GFP*) for cluster 6. Tg(*gcga:GFP*) and Tg(*trpa1b:GFP*) reporters and anti-PYY and anti-CCK antibodies were already available [89–92], but we developed novel *ghrl* and *ngn3* promoter-driven QF2 reagents (S5 Fig and Methods). These reagents leverage the QF2-QUAS binary expression system, enabling us to combine our QF2 reporters with different QUAS tools [93,94]. As expected, our *ghrl* reporter co-labels with the *neurod1* reporter and with cells that stain with anti-Ghrelin antibody (S5C-S5L Fig). Our *ngn3* reporter likewise co-labels with the *neurod1* reporter and labels a similar number of cells in the intestine as previously reported lines generated with the same promoter (S5M-S5N Fig) [95].

**Fig 3.**
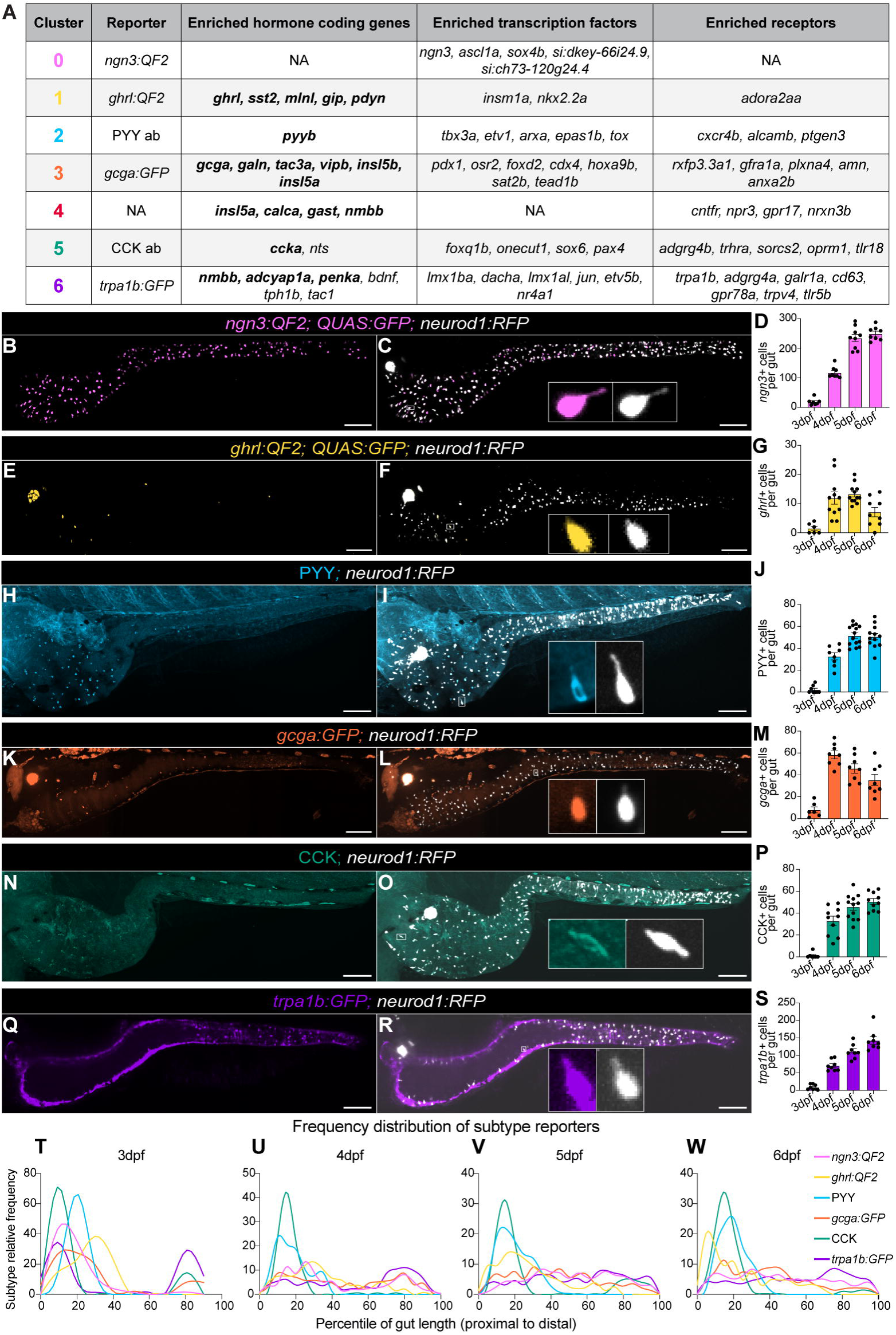
*In vivo* imaging of EEC subtypes shows distinct spatial and temporal patterns during larval development. **(A)** Summary of the hormones, transcription factors, and receptors significantly enriched in each EEC cluster and the reporter used to characterize that cluster. Bolding indicates genes from which we detected peptides. **(B-S)** Representative images from 6 days post fertilization (dpf) fish show subtype distribution and overlap with pan-EEC reporter *neurod1.* Individual cells are shown in inset. Quantitation of EEC subtype cell numbers from 3-6 dpf fish are also shown with each dot representing an individual fish. All scale bars are 100 μm. **(T)** 3dpf, **(U)** 4dpf, **(V)** 5dpf, and **(W)** 6dpf subtype localization data shown as frequency distributions of cells’ locations along the length of the gut divided into percentiles from proximal to distal. Distributions are smoothened for visualization.

To understand the dynamics of these EEC subtype reporters during intestinal development, we crossed each of them to the pan-EEC reporter *neurod1:RFP* [26] and imaged them daily from 3 to 6 days post fertilization (dpf) (S6, S7 Figs). Cell counts from each day and representative images at 6dpf are shown (Fig 3B-3S). Subtype reporters showed a variety of prevalence and regional distributions (Fig 3T-3W), which may contribute to their sub-specialized functions. *ghrl+,* PYY+, and CCK+ cells – labeling subtype clusters 1, 2, and 5, respectively – were all concentrated proximally within the intestine. Mammalian orthologs *Ghrl/GHRL* and *Cck/CCK* also show a proximal skew in the small intestine, but *Pyy/PYY* is more prevalent in the distal small intestine and colon in mammals [1,96]. *gcga+* cells from subtype cluster 3 peaked in the midgut, consistent with the enrichment of the transcription factor *osr2* in this cluster [71]. In accordance with mammalian literature, *trpa1b*+ cells labeling subtype cluster 6 were the most prevalent subtype and were relatively evenly distributed along the length of the intestine [97]. Overall, the subtype reporters appeared as early as 3dpf but became more widely expressed at 4dpf, in alignment with *neurod1* reporter activity, growth of the intestine, and *in situ* hybridization data [29,30]. After 4dpf, EEC subtype numbers increased with age except for *gcga+* and *ghrl*+ cells, which showed a trend towards decreasing after their initial burst of cell number at 4dpf.

### Ablation of *ngn3+* cells leads to reduction in EECs limited to select subtypes

We predicted that cells in scRNA-seq cluster 0 were EEC progenitors based on their enrichment with *ngn3* and other EEC progenitor transcription factors and their early position on the pseudotime-predicted differentiation trajectory (Fig 1G, 1H). As cluster 0 made up a relatively small population on the UMAP (Fig 1B), we were surprised to see our *ngn3* reporter label so many EECs (Fig 3B-3D). As the mouse ortholog, *Ngn3*, is known to have transient expression in EEC progenitors [98], we speculate that the perdurance of both QF2 and GFP is contributing to a long-lasting labeling of cells that transiently expressed *ngn3* in the past. The higher than expected number of *ngn3*+ cells therefore suggested that many EECs outside of cluster 0 at one time expressed *ngn3*.

To test this hypothesis, we used a combination of reporters to genetically induce Diphtheria toxin subunit A (DTA) expression, and thus cell death, in any *ngn3*-expressing cell in the intestine (Fig 4A). Specifically, we leveraged a reporter that uses *gata5* regulatory sequences to drive intestinal epithelial expression of a loxP-flanked mCherry upstream of DTA [27]. When we combined this reporter with our *ngn3:QF2* and *QUAS:Cre* alleles, the mCherry is recombined out in *ngn3*+ cells, leading to expression of DTA and cell death, which we operationally call “*ngn3+* cell ablation.” This strategy affects cluster 0, which has enriched *ngn3* expression, and effectively prevents the emergence of any further mature subtypes that ever pass through a *ngn3*+ state. We evaluated the total numbers and regional distribution of *neurod1*+ cells (Fig 4B) in this *ngn3+* cell ablation model and found that roughly 50% of all EECs are lost with the effect being more pronounced in the proximal gut (Fig 4C). This suggests that approximately half of all EECs pass through a *ngn3+* cell state, showing that, in contrast to mammalian systems, a *ngn3+* cell state is not required for development and differentiation of all EECs in zebrafish [10].

**Fig 4.**
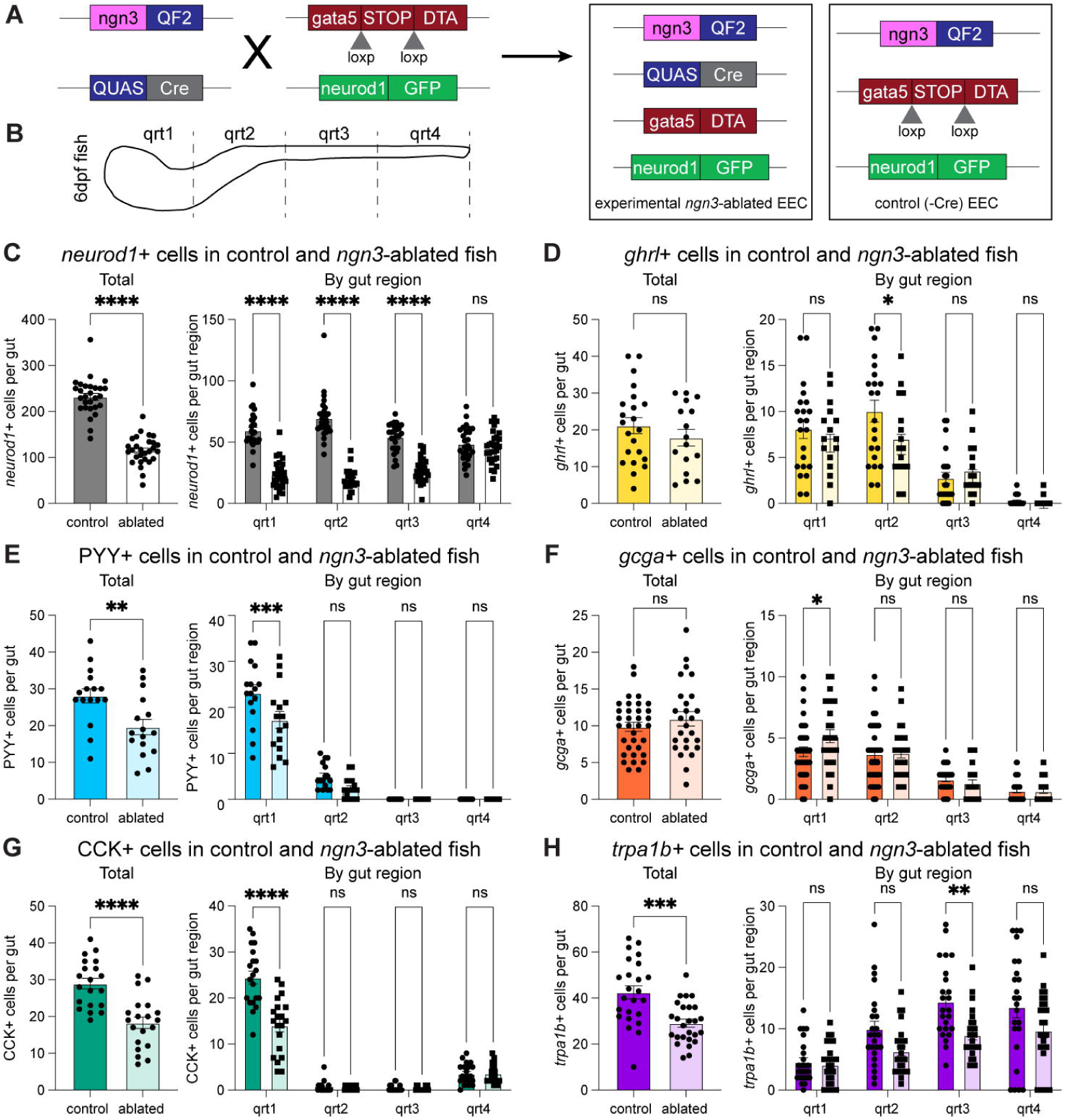
Selective ablation of *ngn3*+ cells in the intestine reduced total EECs and several EEC subtypes. **(A)** Schematic of *ngn3*+ cell ablation. **(B)** Schematic of regional scoring system. Total and regional counts of **(C)** *neurod1*+ cells, **(D)** *ghrl*+ cells, **(E)** PYY+ cells, **(F)** *gcga*+ cells, **(G)** CCK+ cells, and **(H)** *trpa1b*+ cells in control and *ngn3* ablated fish are shown. Each dot represents a 6 day post fertilization fish. Statistical significance was calculated by unpaired *t* test for total cell numbers and by two-way ANOVA for regional analysis. Significance annotations are as follows: ns (p>0.05), * (p<0.05), ** (p<0.01), *** (p<0.001), **** (p<0.0001).

We next combined our subtype reporters with our ablation constructs to ask if certain EEC subtypes were particularly affected by *ngn3*+ cell ablation. Because we could not combine the *ngn3:QF2* and *ghrl:QF2* reporters, we generated a *ngn3:Cre* line to test the effects of *ngn3*+ cell ablation on *ghrl*+ cells. We confirmed that *ngn3+* cell ablation induced with the *ngn3:Cre* reporter produced similar effects on *neurod1*+ cell numbers as the *ngn3:QF2; QUAS:Cre* reporter (S8 Fig). When looking across EEC subtype populations, we found that PYY+, CCK+, and *trpa1b+* cells were all reduced by roughly 30% while *gcga+* and *ghrl*+ cells were unaffected by *ngn3*+ cell ablation (Fig 4D-4H). Consistent with our reporter characterization (Fig 3), reductions in EEC subtypes were observed in the region of the gut where they were most prevalent – proximally for PYY+ and CCK+ cells and distally for *trpa1b*+ cells. This suggests that a portion of PYY+, CCK+, and *trpa1b+* subtypes pass through a *ngn3+* cell state during their differentiation.

### Ablation of *ghrl+* cells leads to reduction in EECs in subtypes distinct from *ngn3*+ ablation

The persistence of *ghrl+* cells despite *ngn3+* cell ablation is reminiscent of mammalian data that shows *Ghrl* expression and *Ghrl*+ cell numbers were maintained or increased upon deletion of a host of transcription factors important for EEC differentiation like *Nkx2.2a* and *Pax4* [19,64,65,99–102]. In addition, our pseudotime analysis predicted a differentiation trajectory moving through *ghrl*+ cells, strikingly similar to several mammalian scRNA-seq datasets that suggested a *Ghrl*+ EEC progenitor population [23–25]. Older lineage tracing data also supports a role for *Ghrl*+ cells as progenitors of other endocrine populations in the stomach and pancreatic islet [22]. However, the hypothesis that *Ghrl*+ cells in the intestine give rise to other EEC subtype populations remained untested.

Motivated by the parallels between our data and published studies in mammals described above, we leveraged our toolkit to test if any EEC subtypes pass through a *ghrl*+ cell state. Using a similar genetic approach as above, we selectively killed any *ghrl*-expressing cell in the intestinal epithelium, which we operationally define as “*ghrl+* cell ablation” (Fig 5A). Despite *ghrl*+ cell numbers peaking at approximately 20 cells per animal during larval development (Fig 3G), we found *ghrl*+ cell ablation resulted in a loss of roughly 40 EECs, indicating other EEC subtypes might be affected (Fig 5B). Indeed, we found a 30% reduction in PYY+ and *gcga*+ EECs in *ghrl*-ablated fish while CCK+ and *trpa1b*+ cells remained unaffected (Fig 5C-5F). Notably, this is distinct from the *ngn3+* ablation result where PYY+, CCK+, and *trpa1b*+ cells were reduced and *ghrl+* and *gcga+* cells were maintained (Fig 4). These data suggest a *ngn3*- independent role for *ghrl*+ cells in the differentiation of PYY+ and *gcga*+ EEC subtypes. This could be explained by PYY+ and *gcga*+ cells passing through a *ghrl*+ cell state or by the loss of a secreted factor from *ghrl*+ cells that promotes differentiation of these subtypes.

**Fig 5.**
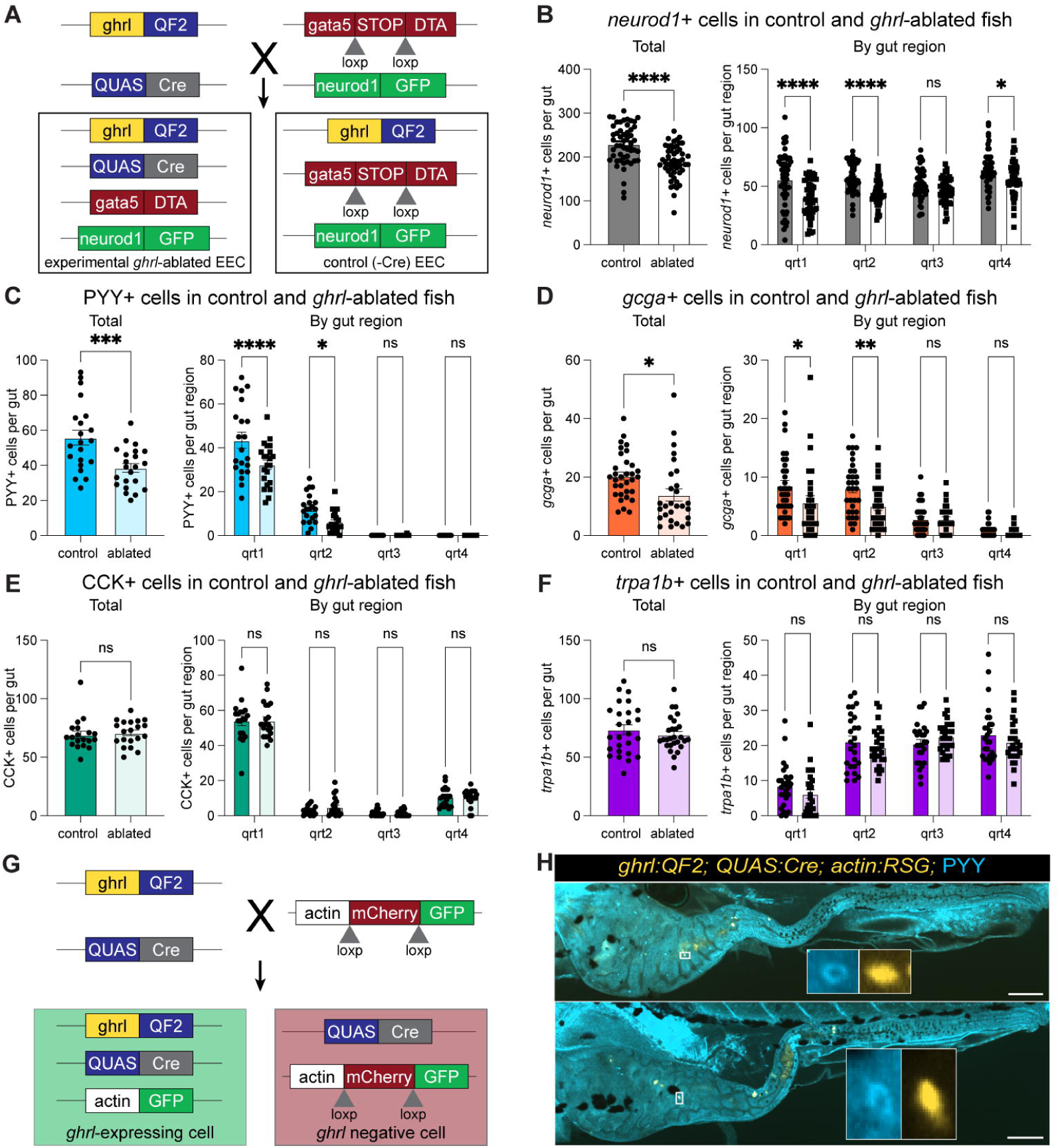
Selective ablation of *ghrl*+ cells suggests lineage relationships between *ghrl*+ cells and other EEC subtypes. **(A)** Schematic of *ghrl*+ cell ablation. Total and regional counts for **(B)** *neurod1*+ cells, **(C)** PYY+ cells, **(D)** *gcga*+ cells, **(E)** CCK+ cells, and **(F)** *trpa1b*+ cells in control and *ghrl* ablated fish are shown. Each dot represents a 6 day post fertilization fish. **(G)** Schematic of *ghrl+* cell lineage tracing. **(H)** Representative images of *ghrl* lineage tracing with PYY staining showing rare double positive cells. Scale bars are 100 μm. Statistical significance was calculated by unpaired *t* test for total cell numbers and by two-way ANOVA for regional analysis. Significance annotations are as follows: ns (p>0.05), * (p<0.05), ** (p<0.01), *** (p<0.001), **** (p<0.0001).

To distinguish these possibilities, we first used lineage tracing tools to indelibly label a cell with GFP if it has ever expressed *ghrl* (Fig 5G). We then performed PYY staining and found rare occurrences of double positive cells, indicating that some PYY+ cells have indeed passed through a *ghrl*+ cell state (Fig 5H). This suggests that the impact of *ghrl+* cell ablation on PYY+ cells is at least in part cell-autonomous. Due to overlapping fluorophores, we were not able to do a similar study with *gcga*+ cells.

Given the small number of *ghrl*-derived PYY+ cells in our lineage tracing analysis, we hypothesized that *ghrl*+ cells could also be regulating other EEC subtypes non-cell autonomously through secreted products. As ghrelin peptides were prevalent in our peptidomics dataset (Fig 2D) and have been shown to stimulate growth of intestinal cells [103,104], we tested if loss of the *ghrl* gene impacted EEC development. We used CRISPR/Cas9 to generate a *ghrl* mutant and evaluated the effects on *neurod1*+ EECs. We found that EEC numbers were equivalent across mutant and wildtype fish (S8 Fig), suggesting that loss of *ghrl* does not overtly impact EEC subtype differentiation.

## Discussion

Here we reported seven distinct EEC populations in zebrafish and characterized reporters for six of them. Our characterization of EEC subtype reporters is consistent with previously published larval zebrafish *in situ* data, building confidence in our proposed EEC subtype identities [28,30]. These studies agree with our data that first onset of these genes is around 3dpf (72hpf) with expression increasing by 4dpf [28,30]. While PYY and CCK antibody staining cells were rare at 3dpf in our study, *pyyb* and *ccka in situ* hybridization data showed a greater number of positive cells [30], suggesting that protein formation lags slightly behind transcript presence, as might be anticipated. Despite this slight difference, the regional localization of these cells across all timepoints and the number of positive cells at 4dpf were consistent between our observations and published studies [28,30]. Intriguingly, *in situ* hybridization analysis of genes that we reported were expressed in the same EEC subtype showed the same prevalence and regional patterns as each other, consistent with them being present in the same cells. For example, EEC subtype cluster 1 in our scRNA-seq dataset expressed *ghrl, gip, mlnl,* and *sst2* and was labeled in this study with a *ghrl* reporter. Both our *ghrl* reporter and published *in situ* data for *ghrl, gip, mlnl,* and *sst2* labeled <10 cells in the proximal gut at 4dpf [30].

Furthermore, previous studies showed *ghrl* and *mlnl in situ* probes overlap, as would be predicted from our data [30]. Also our identification of EEC cluster 6 as enterochromaffin cells is consistent with previous reports of colocalization of enterochromaffin markers *trpa1b*, *tph1b,* and serotonin in a subset of zebrafish EECs that communicate with the nervous system [27].

This consistency builds confidence in our findings and suggests that our observations from EECs at 6dpf may also be true at earlier timepoints.

We show larval and adult EECs have remarkably consistent subtype signatures (Figs 1D, 1E, 1G S2D-S2F), suggesting that EEC subtype programing is established in unfed larvae and persists through adulthood. This indicates that the developmental mechanisms underlying EEC subtype programs are relatively resilient to environmental and ontogenetic changes and that findings in one age group may be generalizable to another. Similar studies across ages in other animals are limited. One scRNA-seq study of the human intestine looked at EECs from adult, pediatric, and fetal tissue samples. Although a comparison across sample age was limited by 90% of the EECs coming from fetal tissue, the cells did not clearly separate by age. Instead, as in our study, clustering was largely driven by hormone expression [74]. Additional studies of human fetal EECs have likewise shown hormone expression patterns similar to those reported from adult tissue [51,105,106]. This suggests that the establishment of EEC subtype programs during early development that endure into adult stages may be a common feature of EECs in fishes as well as mammals.

The repertoire of hormone coding genes expressed in EECs is very similar between zebrafish and mammals (Fig 1F), despite over 400 million years having elapsed since their last common ancestor [107]. Almost all the EEC hormone coding genes we report in our zebrafish dataset are found in mammalian EECs, with the exception of *pdyn. PDYN* is present in mammals but has not yet been reported in EECs. Conversely, four commonly reported mammalian EEC hormones were not found in our scRNA-seq dataset. Two of them – *SECRETIN* and *PPY*– do not appear to have orthologs in zebrafish. *IAPP* has an orthologous novel gene that has not been well described in zebrafish and *NPY* similarly has a zebrafish ortholog, but has largely been characterized in the brain [108,109].

We went beyond hormone gene expression to report the first comprehensive catalog of zebrafish EEC peptides, demonstrating that zebrafish EECs make highly processed and modified peptide hormones. We report 425 unique peptides derived from 18 EEC hormone coding genes (Fig 2B). Of note, our liquid chromatography and mass spectrometry settings may have slightly biased the peptides we were able to detect here. For example, as in many standard protocols, singly charged peptides were not selected for fragmentation, meaning that we would not identify many Leu and Met enkephalin peptides that might be derived from *proenkephalin a* (*penka*) or *pdyn*. Furthermore, the complexity of peptidomic samples, including the high charge states and complex fragmentation, complicate the detection of longer peptides, although peptides up to 60 amino acids long have been reported using similar techniques [110,111]. Finally, while we analyzed both larval and adult samples, the total cell content in adult samples was almost twice that of larval samples (50,000 versus 30,000 cells) due to limitations in sorting and collecting EECs from larvae. Thus, while peptides were detected at higher levels and with greater frequency in adult samples, we cannot conclude that they were absent in larval samples as they may have been below the limit of detection. For this reason, we did not perform direct comparisons between larval and adult samples here.

We hope the zebrafish EEC peptides we report here will be leveraged for further discovery related to EEC hormone endocrinology, and we refer the reader to the supplemental materials for a complete catalog of the detected peptides (see S3 Fig and S2 Table). There are exciting possibilities for discovery related to the peptides we report here that show homology to known human hormones. Historically, studies of homologous peptide hormones in non-mammalian models have provided critical insights into hormone function and even informed therapeutic design. For example, the first identification of GLP-1 occurred in anglerfish [112], and Exendin-4, a peptide hormone produced by gila monster lizards that is analogous to human GLP-1, contributed to the development of long-acting GLP-1 analogs now broadly used for treatment of diabetes and obesity in humans [113–115]. We detected several zebrafish peptides that map to the regions of *gcga* and *gcgb* that align with the GLP-1 encoding portion of human *GCG* (S3B Fig). The arginine 36 residue in GLP-1 is commonly amidated in mammals [116,117], as is seen with one of the *gcgb*-derived peptides detected in zebrafish. However, there are also notable differences between zebrafish *gcga/gcgb* and human *GCG* and their corresponding peptides.

Firstly, *gcgb* and isoform 2 of *gcga* appear to have lost the GLP-2 coding region, a phenomenon also reported in other fishes [118,119]. Additionally, the GLP-1 region in zebrafish *gcga* lacks the highly conserved dibasic PCSK1/2 cleavage site at its C-terminus [120], suggesting alternative processing of this peptide, perhaps at the dibasic RR site immediately upstream of the GLP-2 region. This would result in a 45 amino acid peptide, which would be more challenging to detect and might explain why, although there are many detected peptides aligning to the GLP-1 region in zebrafish, there is not one that perfectly overlaps with the predicted GLP-1 sequence. These nuances in peptide structure and sequence could provide valuable insight into the evolution, function, and regulation of these hormones. Detected peptides aligning to regions annotated in humans as coding for peptide hormones like PACAP, glucagon, glicentin related polypeptide, calcitonin, katacalcin, galanin, INSL5 A and B chain, motilin, neuromedin B, PENK, Leu and Met enkephalins, synenkephalin, neuronostatin, and somatostatin could likewise be exciting candidates for future studies into EEC peptide hormone biology. There are also opportunities to potentially uncover novel signaling molecules among the peptides that are abundant in zebrafish but align to a region with no annotated function in humans. Indeed, recent studies have illustrated how new technologies and datasets such as this can continue to reveal novel peptide hormones [76,77,121,122]. While these are a few examples, we hope the entire zebrafish EEC peptide catalog will be a resource for future discovery related to EEC hormones (S3 Fig, S2 Table).

We also detected peptides derived from *glucagon b, somatostatin 1.2,* and *insulin* (Figs S3B, S3M, S3P) hormone genes that are commonly thought to be enriched in the pancreatic islet [30,123,124]. While this could be evidence of pancreatic contamination in our sorted EEC samples, those genes do show some limited expression in the joint larval and adult EEC dataset (S4E Fig) and have been reported in zebrafish intestinal bulk RNA datasets before [30,125], suggesting these peptides could be produced in zebrafish EECs.

Despite striking conservation of EEC hormone genes and peptides, the way those hormones are partitioned into EEC subtypes shows notable differences between zebrafish and mammals. Specifically, while *gip* and *nts* are present in zebrafish EECs, *gip* or *nts* expressing cells do not separate into their own clusters as they often do in mammalian datasets, perhaps because they are expressed at relatively lower levels in zebrafish [19,52,53]. Furthermore, while *PYY* and *GCG* are found together in what have traditionally been called “L cells” in mammals, we see their orthologs *pyyb* and *gcga* separate into clusters 2 and 3, respectively, in our dataset. Of note, cluster 2 and 3 are somewhat intermingled in the UMAP plot and have the most overlapping marker profile of any two clusters (Figs 1G, S2D-S2F), suggesting a close relationship between the two populations. Interestingly, scRNA-seq reports from mouse colon, where *Pyy* is more highly expressed, have separated *Pyy* and *Gcg* into separate clusters [96]. Although the zebrafish intestine most closely resembles the mammalian small intestine [126–128], we also observed high expression of *insulin like 5* paralogs *insl5a* and *insl5b* in our dataset, the mammalian homolog of which is mostly restricted to colonic EECs [51,54,96,129]. These findings suggest that some differences in EEC subtype identities between zebrafish and mammals may be due to sub-specialization of intestinal regions in mammals.

Our results indicate that *ngn3* has a smaller role in EEC differentiation in zebrafish than in mouse. *Ngn3* has been shown to be necessary and sufficient for committing secretory progenitors to an endocrine fate in both pancreatic islet endocrine cells and EECs in mammals [10–14,130–132]. Although *ngn3* was shown to be dispensable for islet development in zebrafish, its role in EECs had not been formally tested [29]. We observed a more minor role for *ngn3+* cells in EEC differentiation in zebrafish where only some EEC subtypes – namely *trpa1b+,* CCK+, and PYY+ cells – seem to pass through a *ngn3*+ cell state. This contrasts with our pseudotime data, which predicted that most EEC subtypes, and specifically *ghrl*+, PYY*+,* and *gcga+* populations, would all pass through an EEC progenitor population enriched for *ngn3* expression (Fig 1H). This discrepancy could be because *ngn3* labels only a portion of our putative EEC progenitor population, such that some EEC progenitor cells in cluster 0 are unaffected by *ngn3* ablation and are still able to differentiate into EEC subtypes. Alternatively, the more limited role of *ngn3*+ cells we observed *in vivo* could be because cell intermediates of alternative differentiation trajectories from secretory progenitors to differentiated EEC subtypes might have been poorly captured in our scRNA-seq data. For example, a few secretory progenitor cluster 8 cells cluster with *ghrl*+ cluster 1 cells, raising the possibility that there could be a path from cluster 8 to cluster 1 that does not pass through *ngn3*+ cells in cluster 0. Finally, ablation studies always carry the potential caveat of compensation, in which removing one population makes room for another to at least partially take its place. Despite these methodological limitations, our *in vivo* studies demonstrate that ablation of *ngn3*+ cells only impacts a subset of EEC subtype populations. This partial effect of *ngn3* cell ablation is consistent with published *ngn3 in situ* hybridization data [28] and findings that support *neurod1* as the major fate determining transcription factor in zebrafish EECs [27,31]. By discerning the specific role of *ngn3* in zebrafish EECs, our work helps uncover the processes that govern EEC differentiation across vertebrates, although the specific factors that promote the role of *neurod1* over *ngn3* and what impacts this has on EEC differentiation remain unresolved.

Prior studies in mammals raised the intriguing possibility that *Ghrl-*expressing EECs may have a role in the differentiation of EEC progenitors into mature EEC subtypes, but this had not been directly explored. Reports showed *Ghrl* expression and *Ghrl*+ cell numbers were uniquely resistant to deletion of transcription factors important for EEC differentiation, such as *Nkx2.2* [64–66], *Arx* [100,133], *Pax4* [100,102], *Sox4*, *Tox3*, [19] *Myt1* [19], and *Isl1* [101]. While *Ghrl*+ cell numbers were maintained or often increased across these studies, almost all other EEC subtype markers were reduced. Notably, our data showing *ghrl*+ cells were unaffected by *ngn3*+ cell ablation is consistent with this pattern. Intriguingly, no evidence was found that *Ghrl*+ cells were actively cycling or proliferating [64,102,134]. Altogether, these data support two possible models to explain why *Ghrl*+ cells are uniquely enriched upon loss of various transcription factors important for EEC differentiation: 1) *Ghrl*+ cells are an early-arising mature EEC fate that becomes the default destination for progenitors in the absence of key transcription factors; or 2) *Ghrl*+ cells themselves give rise to other subtypes through the subsequent expression of key transcription factors, and, when those transcription factors are lost, EECs become arrested in a *Ghrl*+ state. Data from different scRNA-seq studies in mammalian EECs are consistent with both models. Computational analysis of time series scRNA-seq data from mouse organoids show that *Ghrl*+ cells are some of the first to arise [19], supporting the first model. However, additional scRNA-seq studies have identified “*Ghrl*+ EEC progenitors.” These cells express progenitor markers, such as transcription factors important for EEC differentiation, but also express *Ghrl*, thought to be a mature EEC marker [23–25]. One of these studies went on to perform computational pseudotime analysis and, as in our study, found a predicted differentiation trajectory moving through a *Ghrl*+ EEC progenitor state [24], supporting the second model. Interestingly, studies reporting *Ghrl*+ EEC progenitors also report *Ghrl* expression in later, more mature EEC populations, suggesting the possibility of a hybrid model with two distinct *Ghrl*+ populations, only one of which serves as a progenitor. Despite the preponderance of suggestive evidence, however, a role for *Ghrl*+ cells in EEC differentiation in the intestine has not, to our knowledge, ever been formally tested.

In this study we show that loss of *ghrl+* cells impacts other EEC subtypes, namely *gcga*+ and PYY+ cells, and that *ghrl*+ and PYY+ cells have a lineage relationship, in line with model 2 proposed above. Lineage relationship between *ghrl*+ and *gcga*+ cells could not be tested here due to lack of an antibody labeling *gcga*+ cells. While we acknowledge that lineage-based approaches have limitations and can confuse lineage dependency and co-expression, these data are supportive of a role for *ghrl*-expressing cells in the differentiation of other subtypes.

Interestingly, while lineage tracing of *ghrl*+ cells revealed rare instances of overlap with PYY, these events were not frequent enough to fully explain the reduction of PYY cells seen in *ghrl* ablated fish. As EECs are known to signal to eachother [135–139], it is possible that loss of the signals secreted from *ghrl*+ cells is the primary driver of reductions in other EEC subtypes upon *ghrl+* cell ablation. We did not see any effect on total EEC number following *ghrl* deletion, however, suggesting that loss of ghrelin signaling is not the cause.

Our findings here together with the existing literature support a new model of zebrafish EEC differentiation (Fig 6). EECs are derived from secretory progenitors and are marked by the expression of *neurod1,* which is required for all EECs in zebrafish [27,31]. Here we have shown that a portion of *trpa1b+,* CCK+, and PYY+ EEC subtypes pass through a *ngn3+* and a portion of *gcga*+ and PYY+ EECs pass through a *ghrl*+ cell state. Both *ngn3+* and *ghrl+* cell ablation had only partial impacts on other EEC subtype populations, indicating *ngn3-* and *ghrl-* independent differentiation pathways are also available. Multiple differentiation pathways may contribute to EEC subtype plasticity or compensation, but it is unclear if there are functional differences between cells of the same subtype that differentiate through varying pathways. We also acknowledge that ablation of *ghrl*+ or *ngn3*+ cells may have fundamentally shifted the transcriptional profile of EEC subtype populations. While we can still evaluate the numbers of cells labeling with each subtype reporter, we do not know if the subtype biology beyond that single reporter is preserved.

**Fig 6.**
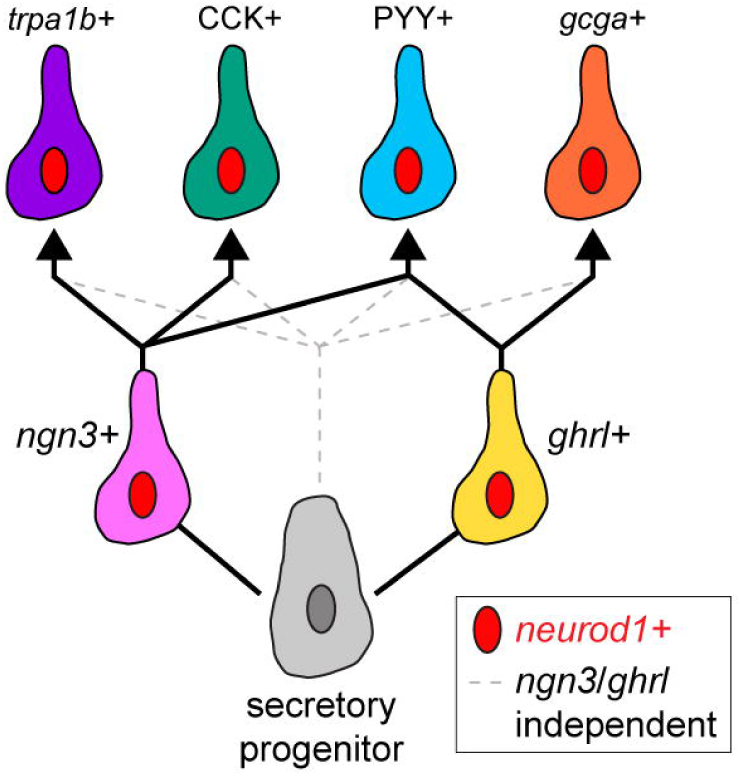
Working model of EEC subtype differentiation in zebrafish.

In conclusion, the tools and datasets reported here can serve as a platform for future hypothesis testing to better understand fundamental EEC subtype and hormone biology. For example, our atlas of endogenous EEC peptides can serve as a starting point for testing their individual endocrine functions and structure-function relationships. The toolkit of EEC subtype reporters also provides a valuable resource for further *in vivo* studies. The modularity of the QF/QUAS transgenic tools deployed here will allow future studies to not only label, but selectively activate or inhibit certain subtypes to understand their physiologic roles. Future work could test the subtype-specific impacts of various challenges such as dietary changes, microbial manipulations, or disease models.

## Materials and Methods

### Ethics statement

All zebrafish studies were approved by the Institutional Animal Care and Use Committee of Duke University (protocol A061-22-03).

### Zebrafish lines and husbandry

Zebrafish stocks were maintained on an Ekkwill (EK) background on a 14-hour/10-hour light/dark cycle at 28.5°C in a recirculating aquaculture system (Pentair). From 6 dpf to 14 dpf, larvae were fed Zeigler AP100 <50-micron larval diet (Pentair, LD50-AQ) twice daily. Beginning at 14 dpf, larvae were fed *Artemia* (Brine Shrimp Direct, BSEACASE) once daily, supplemented with a daily feed of Skretting Gemma Micro 75 (Bio-Oregon, B5676). From 28 dpf, the Gemma Micro 75 diet was replaced with Gemma Micro 300 (Bio-Oregon, B2809). At the onset of breeding age or sexual maturity, adult fish were given a 50/50 mix of Skretting Gemma Micro 500 (Bio-Oregon, B1473) and Skretting Gemma Wean 0.5 (Bio-Oregon, B2818) and one feeding of *Artemia* daily.

For experiments involving zebrafish larvae, adults were bred naturally in system water and fertilized eggs were transferred to 100mm petri dishes containing ∼60 mL of egg water at approximately 6 hours post fertilization. The resulting larvae were raised under a 14 h light/10 h dark cycle in an air incubator at 28°C at a density of 1 larvae/mL water. All the experiments performed in this study ended at 6 dpf unless specifically indicated. As sex determination in zebrafish occurs after 6 dpf, we do not report numbers of male or female fish in each experiment. The following engineered zebrafish lines were used in this study: *TgBAC(cldn15la:EGFP)^pd1034^* [140], *Tg(−5kbneurod1:TagRFP)^w69^*[141], *TgBAC(trpa1b:EGFP)^a129^*[92], *TgBAC(gata5:loxp-mcherry-stop-loxp-DTA)^pd315^*[27], *TgBAC(neurod1:EGFP)^nl1^* [142], *Tg(gcga:EGFP)^ia1^*[143], Tg(β*-actin2:loxP-mCherry-loxP-GFP*)^pd31^ [144], *Tg(QUAS:GFP; cryaa:mCherry*)^pd1199^ [145], *Tg(ngn3:QF2; cmcl2:GFP)*^rdu107^ (generated in this study), *Tg(ghrl:QF2; cmcl2:GFP)*^rdu108^ (generated in this study), *Tg(QUAS:cre; cryaa:mCherry)*^rdu109^ (generated in this study), *Tg(ngn3:Cre; cmcl2:GFP)*^rdu110^ (generated in this study), and *ghrl* ^rdu111^ (generated in this study).

### Construction of zebrafish lines

The Gateway Tol2 cloning approach was used to generate *Tg(ngn3:QF2; cmcl2:GFP), Tg(ghrl:QF2; cmcl2:GFP), Tg(QUAS:cre; cryaa:mCherry),* and *Tg(ngn3:Cre; cmcl2:GFP)* plasmids [146]. The 2114 base pair *ngn3* promoter was previously reported [95] and generously shared by Dr. Olov Andersson (Karolinska Institutet) and cloned into a p5E-MCS vector [146] using TAKARA In-Fusion Snap Assembly master mix (product #638947) with HindIII-linearization of the backbone and amplification of the insert with primers CGGTATCGATAAGCTCCGCGGCCGCCGTACTCGA and ATTCGATATCAAGCTCGGCGCGCCCACCCTTTCTGTA. The pME-QF2 [94], p5E-QUAS [147], p3E-polyA [146], pDestTol2CG2 [146], and pDESTtol2pACrymCherry [148] plasmids were previously reported and obtained from Addgene. The pME-Cre plasmid was previously reported [149] and generously shared by Dr. Mark Cronan. The 647 base pair *ghrl* promoter was amplified from purified (Wizard Genomic DNA purification kit A1120) EK DNA using primers cttaagcttTTCAGAATTCATATCAGATCACAGACACT, cttccgcggCACCTTAGTCTTAATTCTTTGCTACATAC. It was then cloned into the p5E-MCS [146] via digestion of insert and backbone with HindIII (R3104S) and SacII (R01575S) followed by phosphatase treatment (M0289S) and ligation (M0202S). The appropriate p5E, pME, and p3E entry vectors were cloned into destination vectors with either a green heart (*cmcl2:GFP;* pDestTol2CG2) or red eye (*cryaa:mCherry*; pDESTtol2pACrymCherry) as listed above using an LR-Clonase [12538-120) reaction.

The *ghrl* mutant line was generated using CRISPR-Cas9. The guide RNAs (gRNAs) were designed using the “CRISPRscan” tool (www.crisprscan.org/) [150] and purchased fully synthesized from Integrated DNA Technologies. At the one-cell stage, EK strain zebrafish embryos were injected with 1 to 2 nl of a cocktail consisting of Cas9 protein [1000 ng/μl), gRNA [150 ng/μl), and 0.05% phenol red. Injected embryos were screened for mutagenesis with forward primer CACAGTGACAGTTGTAGACTTTAATGCTAAT and reverse primers GTCTCTAAGAAGATTCTCCAGAAGATTCTGA (mutant), AAACATGCTGCTGGCACGGCA (wildtype). The mutations were further determined through Sanger sequencing of the region encompassing the gRNA targeting sites. 10 F0 injected fish were out-crossed to wildtype fish and progeny were screened to test if mutations were transmissible. F1s from F0 founders were then raised to adulthood and sequenced. Two F1s were found to have identical lesions – the deletion started after the first nucleotide in the first exon and continued through the first 17 nucleotides of the third exon – and were bred to generate a stable line. F2s and F3s from this line were used in the reported experiments and were given the allele name *rdu111*. When testing *ghrl* mutants, genotyping of the *mttp* locus was used as a control to verify there was viable DNA in samples using forward primer AGAGACGGTGTCCAAGCAGG and GCTCAAAGACTTTCTTGC with an expected band of 137 base pairs [151].

### mVISTA alignment

To identify conserved regions of the *ghrl* promoter across *Danio* species, we exported the *ghrl* sequence and annotation from ensembl.org [152]. We blasted the *D. rerio ghrl* amino acid sequence against other *Danio* species and extracted 5 kb upstream and downstream of the aligned region for each species. We then put each of these sequences into the mVISTA alignment [153,154], using the *D. rerio* sequence and annotation as the reference. We identified and amplified a 647 base pair region upstream of the *ghrl* transcriptional start site that was highly conserved across a *Danio* species (S5B Fig).

### Larval immunofluorescence and live imaging

All larval imaging was performed on Andor Dragonfly Spinning Disc Confocal plus with a 20x lens in the Duke Light Microscopy Core Facility. For live imaging experiments, zebrafish larvae were anesthetized with Tricaine and mounted in 1% low-melt agarose. For larvae imaged at 3 and 4dpf, the yolk was mechanically removed with fine watchmaker forceps prior to mounting.

Whole mount immunofluorescence staining was performed as previously described [26]. Larvae were euthanized with 5x tricaine and roughly 20 larvae per condition were transferred to a 1.5mL tube. Excess media was removed and 1 mL of chilled 4% paraformaldehyde was added. Larvae were incubated in fixative overnight (>16 hr) at 4°C. Samples were then washed 2x 5 minutes in PBS. For samples of 3 or 4dpf larvae, yolk was then mechanically removed using fine-needle forceps before dehydrating in 50% methanol / 50% PBS solution for 5 minutes.

Larvae were then washed 3x in 100% methanol before incubation in 100% methanol at-20°C for at least 2 hours. Larvae were then rehydrated in serial 50% methanol / 50% PBS, 100% PBS washes and permeabilized at room temperature for at least 20 minutes with PBS with 0.1% Tween-20, 1% DMSO, and 0.3% Triton X-100. Larvae were then washed and blocked with 4% bovine serum albumin solution for at least 30 minutes at room temperature before washing and incubating in primary antibody solution for at least 24 hours at 4°C. Primary antibodies in this study were diluted at 1:100 [(rabbit anti-PYY) [90], rabbit anti-CCK [89], rabbit anti-Ghrl (Anaspec 55529)] or 1:500 (chicken anti-GFP (Aves GFP-1010)). Larvae were then washed every 15 minutes for 2 hours before incubation with Alexa Fluor Invitrogen secondary antibodies diluted at 1:250 (488 goat anti-chicken (A11039), 647 goat anti-rabbit (A21244), 568 goat anti-rabbit (A11011)). Larvae were incubated with secondary antibodies overnight (>16 hours) at 4°C, washed every 15 minutes for one hour, and mounted for imaging.

### Image analysis with ImageJ

Images were processed and analyzed in ImageJ [155]. Images were automatedly blinded such that the scorer knew the reporter but not the condition (e.g., ablated or control) of each image. The start and end of the gut was manually marked using the Cell Counter plugin and x, y coordinates were extracted. *neurod1*+ cells were automatedly counted from maximum intensity projections using the Analyze Particles function with minimum object size of 10 pixels^2^. The number and location of cells was automatedly extracted in R and locations were zero-ed using corresponding manual gut markers. To account for overlapping cells being grouped into a single object, the median object size was used to assign a weight to each object. Objects twice as large as the median size given a weight of 2, objects three times as large as the median were given a weight of 3, and so on, with a minimum weight of 1. Weights were summed to count the total number of cells in an image.

*ngn3*+ cells were counted with a similar automated approach. A composite of *neurod1* and *ngn3* channels was used to identify objects using the Analyze Particles function and the fluorescence of each channel was extracted for each identified object. Objects with a ratio of *ngn3:neurod1* signal >2 were counted as *ngn3*+ only, objects with a ratio of *ngn3:neurod1* signal <0.5 were counted as *neurod1*+ only, and all objects with a ratio in between 2 and 0.5 were counted as double positive. The same adjustment for overlapping cells described above was also applied.

Because of increased background of various sources causing complications with automated thresholding and counting, images from *gcga, trp1b,* and *ghrl* reporters along with PYY and CCK stains were manually counted using the Cell Counter plugin. Counts for reporter characterization in Figure 3 were taken for each slice of the captured image while maximum intensity projections were used for ablation image analysis for higher throughput. In all cases, the number and location of each manually counted cell were extracted for further analysis.

In addition to total counts, the location of automatedly or manually counted cells was determined relative to the manually annotated start and end points of the gut. In cases where a single object was given a weight >1 and counted as multiple cells, the x, y coordinates of the original object was used for each. Cells were binned into four evenly divided bins along the length of the intestine for visualization of counts per quarter of the gut in bar plots (see Figs 4, 5, and S7) or into 100 evenly divided bins for visualization of density per percentile of the gut in density plots (see Fig 3 T-W).

### Adult sections

Adult zebrafish 3-4 months post fertilization of either sex expressing *Tg(neurod1:RFP); Tg(ghrl:QF2; cmcl2:GFP); Tg(QUAS:GFP; cryaa:mCherry)* were retrieved from the circulating aquaculture system before morning feeding and transferred to a separate tank to fast before dissection. Several hours later fish were euthanized in 0.090% 2-Phenoxyethanol and dissected to retrieve the intestine. The intestine was fixed in 4% PFA overnight at 4°C then washed x3 the next day in PBS. The intestine was mounted in 4% Low Melt Agarose in cryomolds and cured at 4°C for 1 hour. 200μm cross sections of the tissue were sectioned via Vibratome (Leica VT1000S) and mounted on slides with Vectashield containing DAPI (Vectashield H-1200). Z-stack images were collected on a Zeiss 780 inverted confocal microscope at either 20x or 40x magnification. Image processing was done in FIJI including maximum intensity projections of z-stacks from each channel.

### EEC FACs sorting

All media were made fresh the morning of with the exception of the 6M Guanidine Hydrochloride, for which the same solution was used throughout the two-week collection period. For larval samples, *Tg(neurod1:RFP); TgBAC(cldn15la:GFP)* adult breeders were crossed and embryos were collected at roughly 6 hours post fertilization and maintained in egg water [156] at approximately 1 embryo/ml density. Unfertilized embryos were removed at 1dpf and fish larvae were anesthetized and sorted for presence of both reporters at 5dpf. The GentleMACS disassociator (Miltenyi 130-093-235) was placed in the 4°C room to come to temperature for the follow day’s experiment. At 6dpf, 75 larvae were euthanized in Tricaine and pooled into a single sample with 9 samples being processed concurrently with single positive and non-transgenic controls. Larvae were transferred with minimal media to Gentle MACS C tubes (Miltenyi 130- 096-334) containing 2 mL of freshly made disassociation buffer [10mg/ml cold protease (Sigma P5380-1G), 10 μM rock inhibitor (VWR S1049), 2.5 mg/ml DNase (Sigma DN25) in Dulbecco’s Phosphate Buffered Saline]. Samples were processed 5 times with protocol C_01 on GentleMACS disassociator with 10 minutes of incubation on shaker at 4°C after each round. 10 ml of FACS media [0.1% BSA (Fisher BP1600-100), 10 μM (VWR S1049) in HBSS (Sigma H9394-1L)] was added and the contexts were poured through a 30 micron strainer (Miltenyi 130-098-458) into a fresh 50 mL conical tube. The strainer was washed with an additional 10mL of FACS media and samples were spun down at 250 x *g* for 5 minutes at 4°C, decanted, and resuspended in 1 mL of FACS media. Samples and controls were then transferred to FACS tubes (Corning 352052) with 5uL of 7AAD (Sigma A9400-1MG) to stain dead cells. Cells were then immediately subjected to FACS at the Duke Cancer Institute Flow Cytometry Shared Resource. *neurod1:RFP; cldn15la:GFP* double positive, 7AAD negative cells from each sample were sorted into a single 1.5mL Lobind tube (Sigma Z666505) with 500uL of 6M Guanidine Hydrochloride for a total of roughly 30,000 cells. This constituted a single sample for downstream peptidomic analysis. Nontransgenic and single transgenic controls (pools of 50 fish per genotype) were prepared as above and used for gating and compensation. A total of three larval peptidomic samples were collected on separate days, all within 1 week of each other.

For adult samples, procedures were largely similar. Male and female *Tg(neurod1:RFP); TgBAC(cldn15la:GFP)* adults were euthanized at 18 months post fertilization and their intestines were dissected, cut along the vertical access to open the gut, and placed in epithelial buffer with 10μM rock inhibitor (5.6 mM Na_2_HPO_4_, 8 mM KH_2_HPO_4,_ 96.2 mM NaCl, 1.6 mM KCl, 43.4 mM sucrose, 54.9 mM d-Sorbitol, 10μM rock inhibitor). Sample rocked gently at 4°C for 15 minutes before being transferred to fresh epithelial buffer with rock inhibitor. Samples were shaken at roughly 150 shakes per minute to remove mucus and fecal debris. Intestines were then transferred to gentle MACS C tubes with 2mL of disassociation buffer, as above. Intestines were digested with 3 rounds of protocol C_01 with 10 minutes or rocking at 4°C after each round.

Identical to above, 10mL of FACS media was then added, and contexts were passed through a 30 micron strainer into a fresh 50ml conical tube. The strainer was then washed with an additional 10 mL of FACS media and the contexts were spun down, decanted, and resuspended in 1 mL of FACs media for immediate sorting. *neurod1:RFP; cldn15la:GFP* double positive, 7AAD negative cells from each of nine samples run in parallel were sorted into a single 1.5mL Lobind tube (Sigma Z666505) with 500uL of 6M Guanidine Hydrochloride for a total of roughly 50,000 cells. This constituted a single sample for downstream peptidomic analysis.

Nontransgenic and single transgenic controls (pools of 50 fish per genotype) were prepared as above and used for gating and compensation. A total of three adult peptidomic samples were collected on separate days, all within 1 week of each other. All samples were stored at-80°C until they were transferred to the Duke Proteomics and Metabolomics Core Facility and processed for downstream peptidomics as described below.

### Protein alignments

Protein sequences for *Danio rerio, Carassius auratus, Ictalurus punctatus, Salmo salar, Oncorhynchus mykiss, <u>Aquarana catesbeiana</u>, Gallus gallus, Rattus norvegicus, Mus musculus, Homo sapiens* were downloaded from UniProt [157]. When more than one sequence was available, the sequence designated as canonical in UniProt was used. If there were multiple relevant entries, a representative one was used for ease of visualization. Sequences were aligned using MUSCLE [158] and visualized with JalView [159]. Percent identity settings were used to color residues.

### Single cell RNA sequencing analysis

We imported raw data from previously published adult [49] and larval [48] zebrafish intestinal scRNA-seq datasets into R (version 4.3.1). Using established markers of intestinal secretory cells, we selected just secretory cells from each dataset (Fig S1) and integrated them using the SCT method in Seurat version 5.0.3 [160]. We further subsetted this dataset for just secretory progenitors and EECs and re-integrated and clustered it to generate the dataset shown in Fig 1A and analyzed in this manuscript. We used monocl3 (version 1.3.4) to perform pseudotime analysis where use_partition was set to TRUE and root_cells were set to cluster 7 [57]. The integrated secretory cell dataset and integrated secretory progenitor and EEC datasets are available as.rds files in our supplementary material along with the R script used to analyze the data.

Of note, the zebrafish *gastrin* gene is currently named either *LOC100536965* or *CR556712.1* in dadRer 11 and is annotated in Ensembl as a noncoding lncRNA (ENSDARG00000117418) for unknown reasons. However, multiple phylogeny studies have identified the exact genetic coordinates of *LOC100536965/CR556712.1* as zebrafish *gastrin* [161–163] and peptides we detected derived from this gene align with RefSeq XM_021479754.1, which is a coding prediction. We therefore refer to the gene as *gastrin*. When searching for expression of gastrin in our scRNA-seq dataset, we found expression under the gene name *CR556712.1* in the larval cells, but no expression of either gene name in the adult cells. Searching using either gene name in the complete published adult dataset [49] also did not return any results, suggesting the gene is either not annotated in that dataset, annotated with a different name, or else not expressed. Detection of *gastrin*-derived peptides in our adult peptidomics samples suggests that *gastrin* is indeed expressed in adult EECs and that poor gene annotation may contribute to this discrepancy.

### Statistical analysis

For the scRNA-seq analysis, statistical analyses for determination of the cluster-enriched markers was calculated using the FindConservedMarkers() function of the Seurat package in R with a Wilcoxon rank sum test. For all other experiments, statistical analysis was performed using unpaired *t* test, or one-way or two-way analysis of variance (ANOVA) with Tukey’s multiple comparisons test with GraphPad Prism. *P* < 0.05 was defined as statistically significant.

### LC-MS/MS analysis of peptidome samples

Samples sorted into 250 μL of 6M guanidine/50 mM ammonium bicarbonate were subjected to five rounds of 30sec bath sonication (Branson) at 30% power with cooling in between. Samples were then diluted with 500μL of 0.1% formic acid and centrifuged at 12,000rpm to remove cellular debris. Lysates were loaded directly onto a 10mg Oasis HLB SPE cartridge (Waters), washed with 2x 1mL of 2% acetonitrile/0.1% formic acid, and eluted with 250 μL of 50% acetonitrile/0.1% formic acid. Eluents were speed-vacuumed to dryness. Samples were then resuspended in 50μL of 50 mM ammonium bicarbonate with 10 mM dithiolthreitol and heated at 70C for 30 min, alkylated with 25 mM iodoacetamide for 30 min at room temperature and then spiked with 2 fmol/μL of pre-digested yeast alcohol dehydrogenase (Waters MassPrep standard).

LC/MS/MS was performed using an EvoSep One UPLC coupled to a Thermo Orbitrap Astral high resolution accurate mass tandem mass spectrometer (Thermo). Briefly, each sample loaded EvoTip was eluted onto a 1.5 µm EvoSep 150μm ID x 15cm performance (EvoSep) column using the SPD30 gradient at 55C. Data collection on the Orbitrap Astral mass spectrometer was performed in a data-dependent acquisition (DDA) mode of acquisition with a r=120,000 (@ m/z 200) full MS scan from m/z 300-2000 in the OT with a target AGC value of 300% and max accumulation time of 50ms. Data dependent MS/MS scans in the Astral were performed on charge states 2-7 from m/z110-2000 at a target AGC value of 200% and max accumulation time of 20ms. HCD collision energy setting of 30% was used for all MS2 scans. Data were imported into PEAKS studio. The software was set to 10ppm mass accuracy on MS1 and 0.02Da on MS2 data with no enzyme rules selected. *De novo* assisted searching was allowed. The MS/MS data was searched against a custom *Danio rerio* database along with a common contaminant/spiked protein database (bovine albumin, bovine casein, yeast ADH, etc.), and an equal number of reversed-sequence “decoys” for false discovery rate determination.

Database search parameters included fixed modification on Cys (carbamidomethyl), variable modification on Met (oxidation), C-terminal amide, and N-terminal acetylation and pyroglutamate. We queried our raw data against a custom database of zebrafish proteins from TrEMBL [157] and Ensembl [152]. To account for genomic polymorphisms within the Ekwill zebrafish strain used here, we supplemented the combined TrEMBL/Ensembl database with protein sequences predicted with customProDB [164] from genetic variants observed in previous RNA sequencing from the same strain [165]. Spectral annotation was set at a maximum 1% peptide false discovery rate based on q-value calculations. Manual verification of spectra of interest were confirmed using targeted extraction/selected ion chromatograms with Skyline (University of Washington, MacCoss Laboratory) [166] including considerations for signal to noise, peak shape, mass errors across isotopologues, and retention time relative to database identifications. The mass spectrometry proteomics data have been deposited to the ProteomeXchange Consortium via the PRIDE partner repository with the dataset identifier PXD058654 [167]. In addition, we have shared screenshots of Skyline data for an example of a manually reviewed peptide in S4C-S4D Fig.

### Ghrelin acylation search

To identify acylated ghrelin peptides, we performed a PEAKS search on a custom database just containing the F1QKX9 ghrelin sequences from Uniprot and selected octanoylation and decanoylation as variable posttranslational modifications in addition the modifications in our original search. Spectral matches and chromatogram peaks of the identified peptides are included in the supplemental materials (S4A-S4B Fig).

## Notes

### Competing Interest Statement

The authors have declared no competing interest.

