## Supplementary figures and images for "Intestinal enteroendocrine cell subtype differentiation and hormone production in zebrafish"

### Sup Fig 1

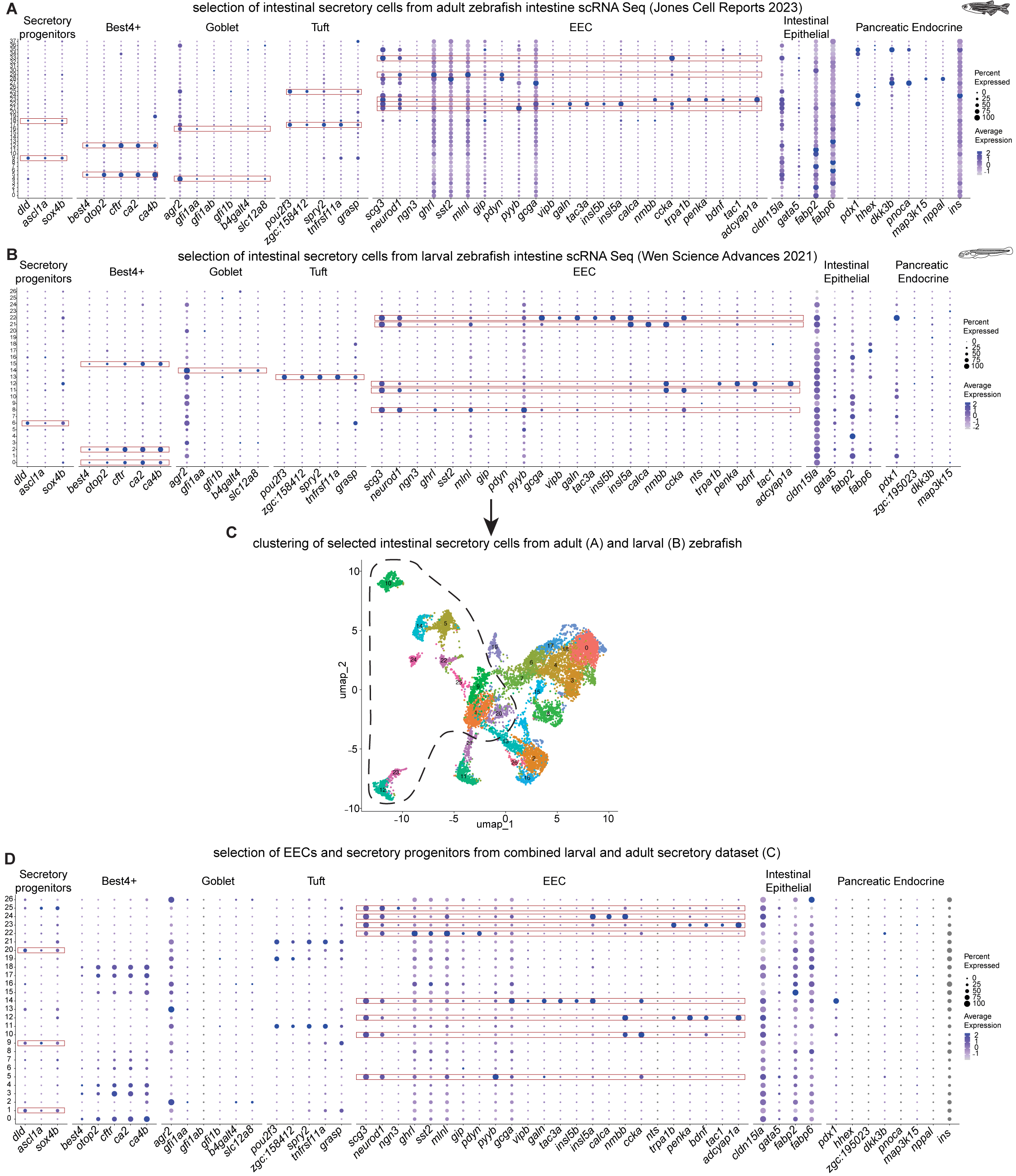

### Sup Fig 2

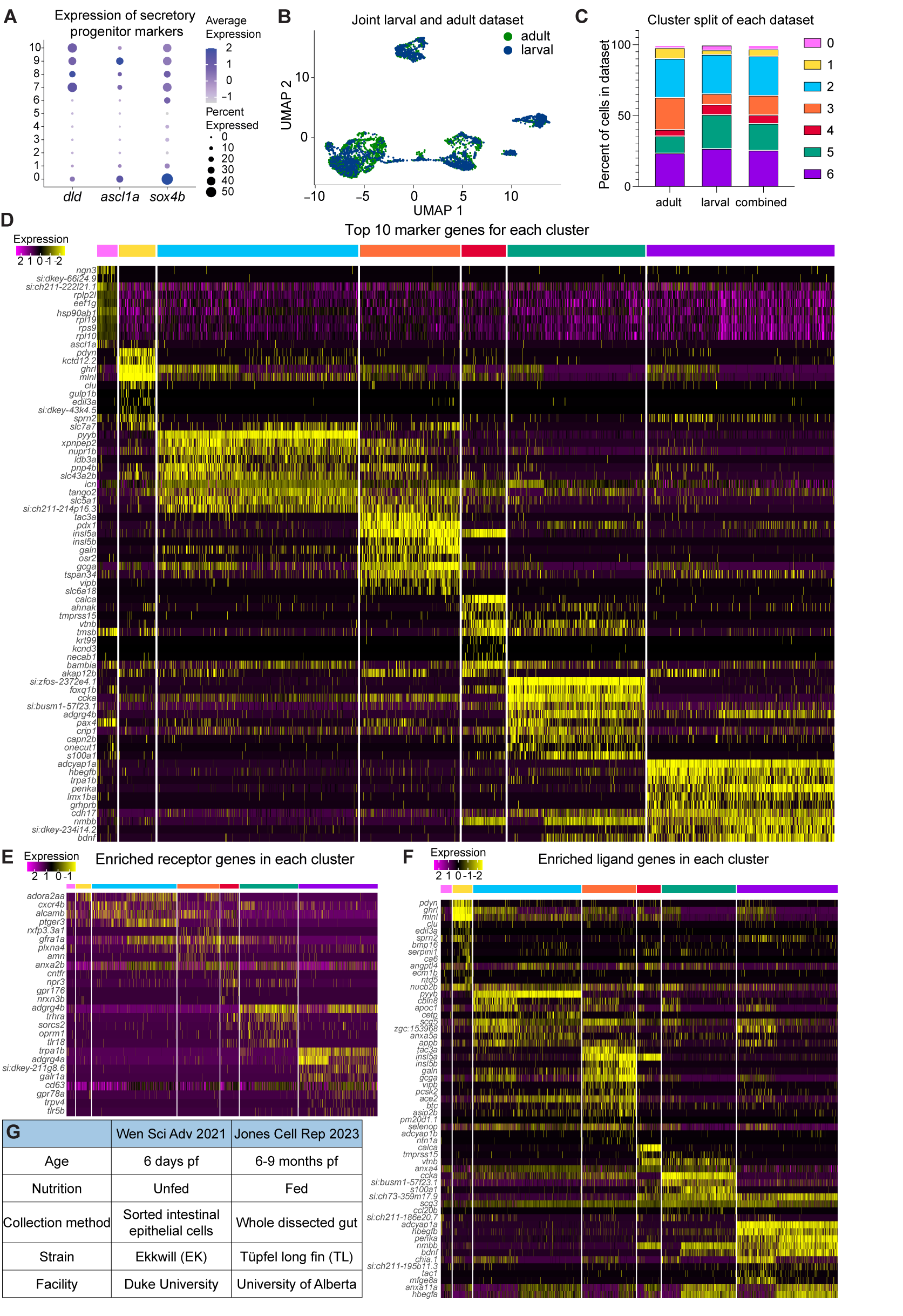

### Sup Fig 4

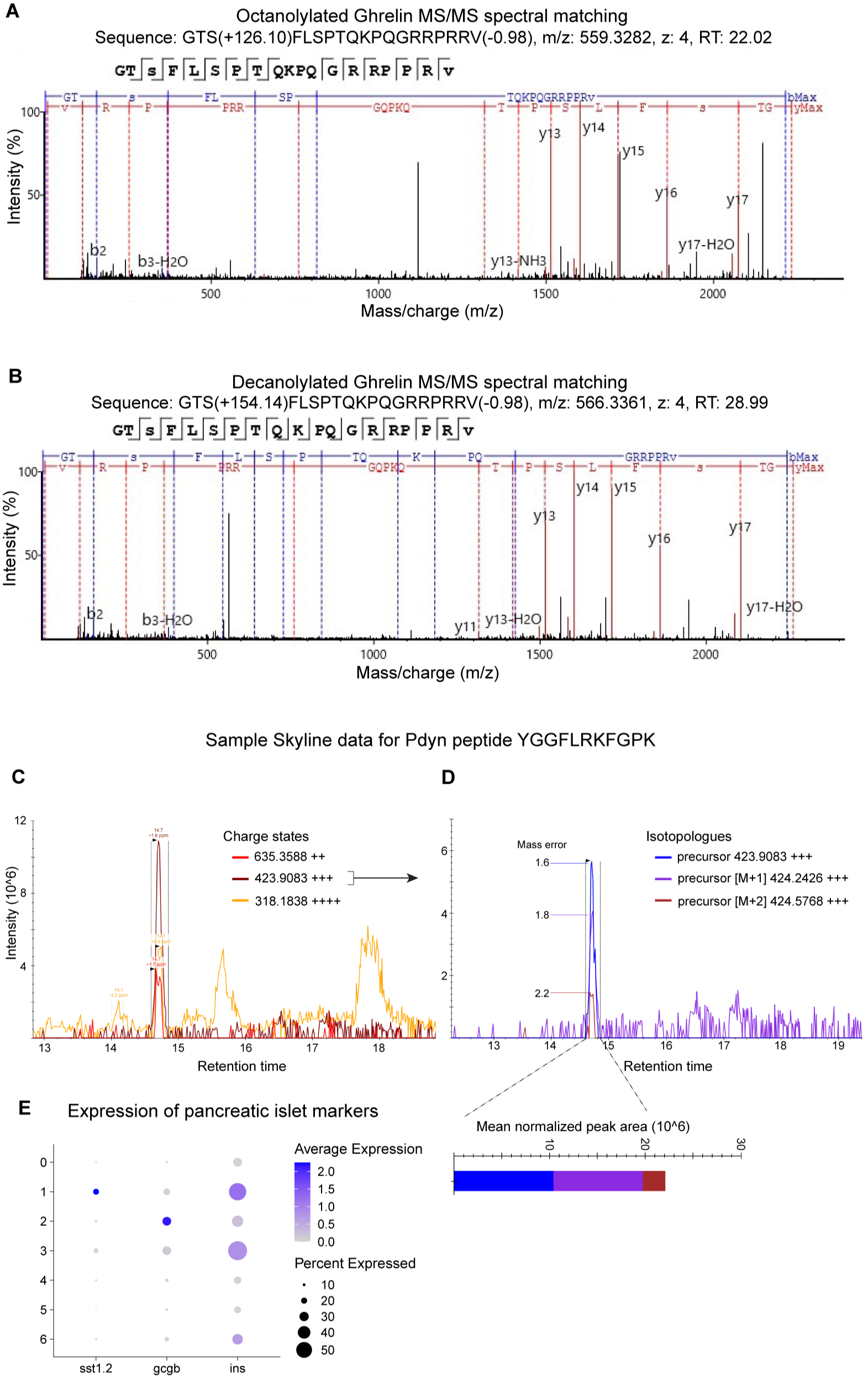

### Sup Fig 5

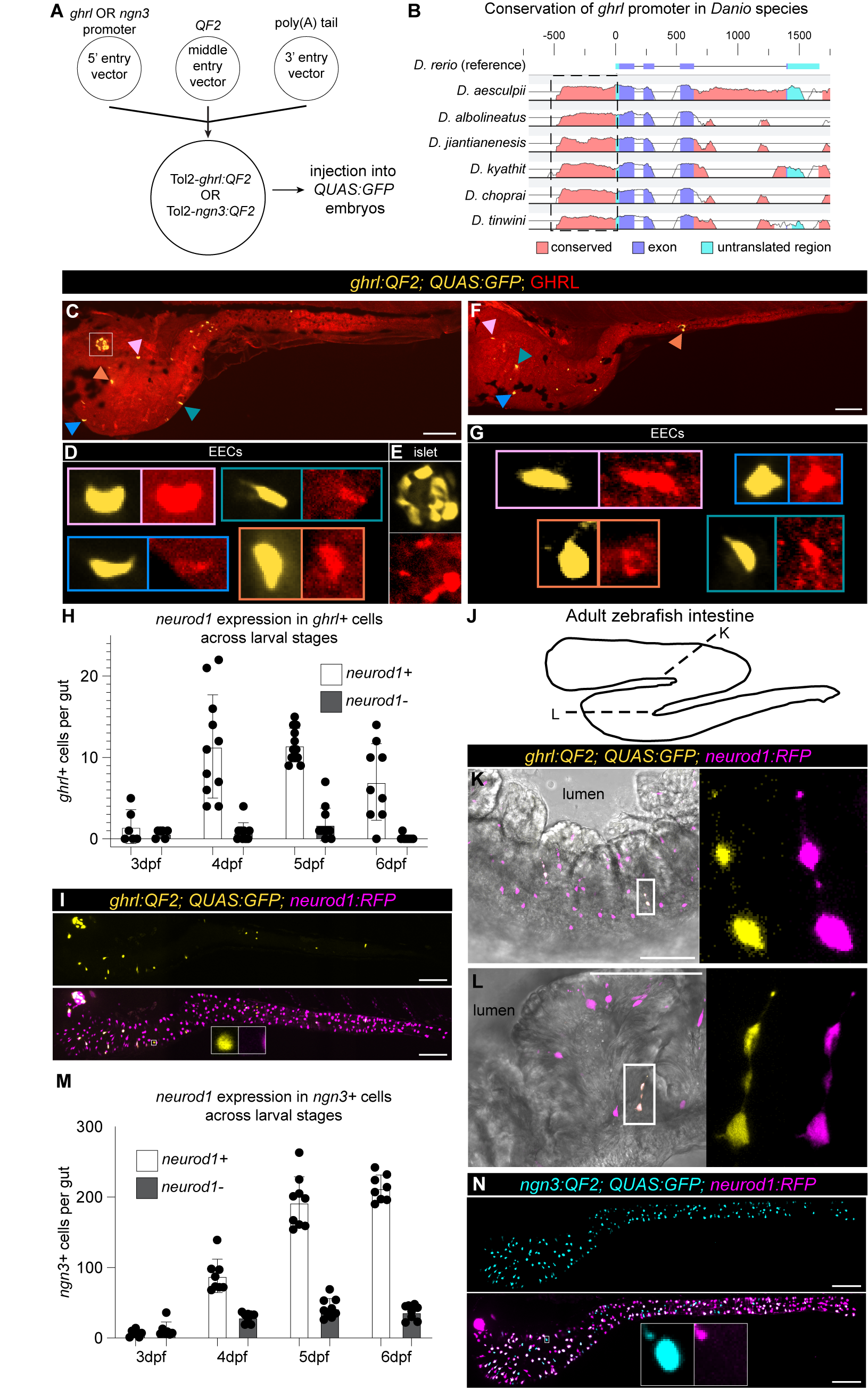

### Sup Fig 6

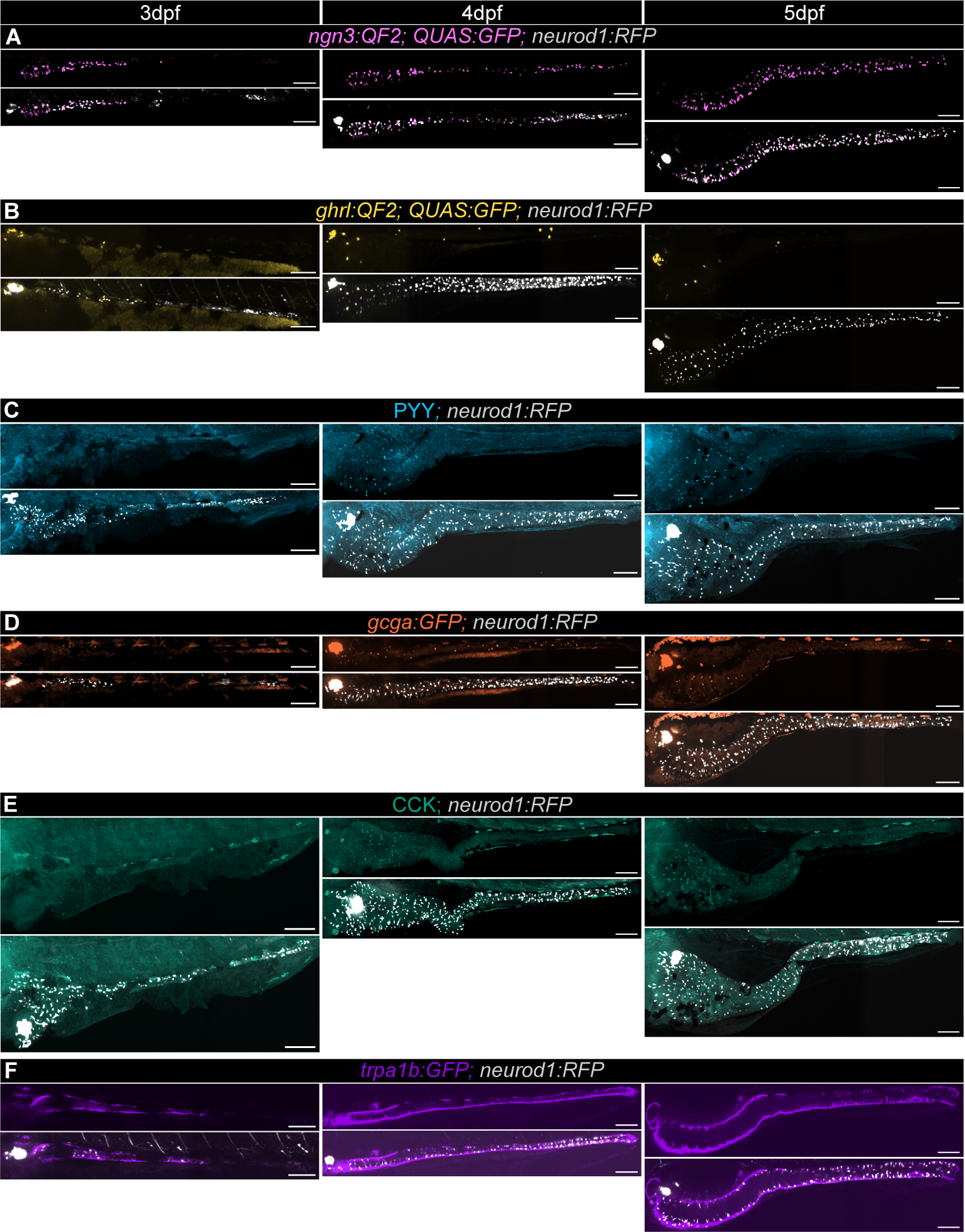

### Sup Fig 7

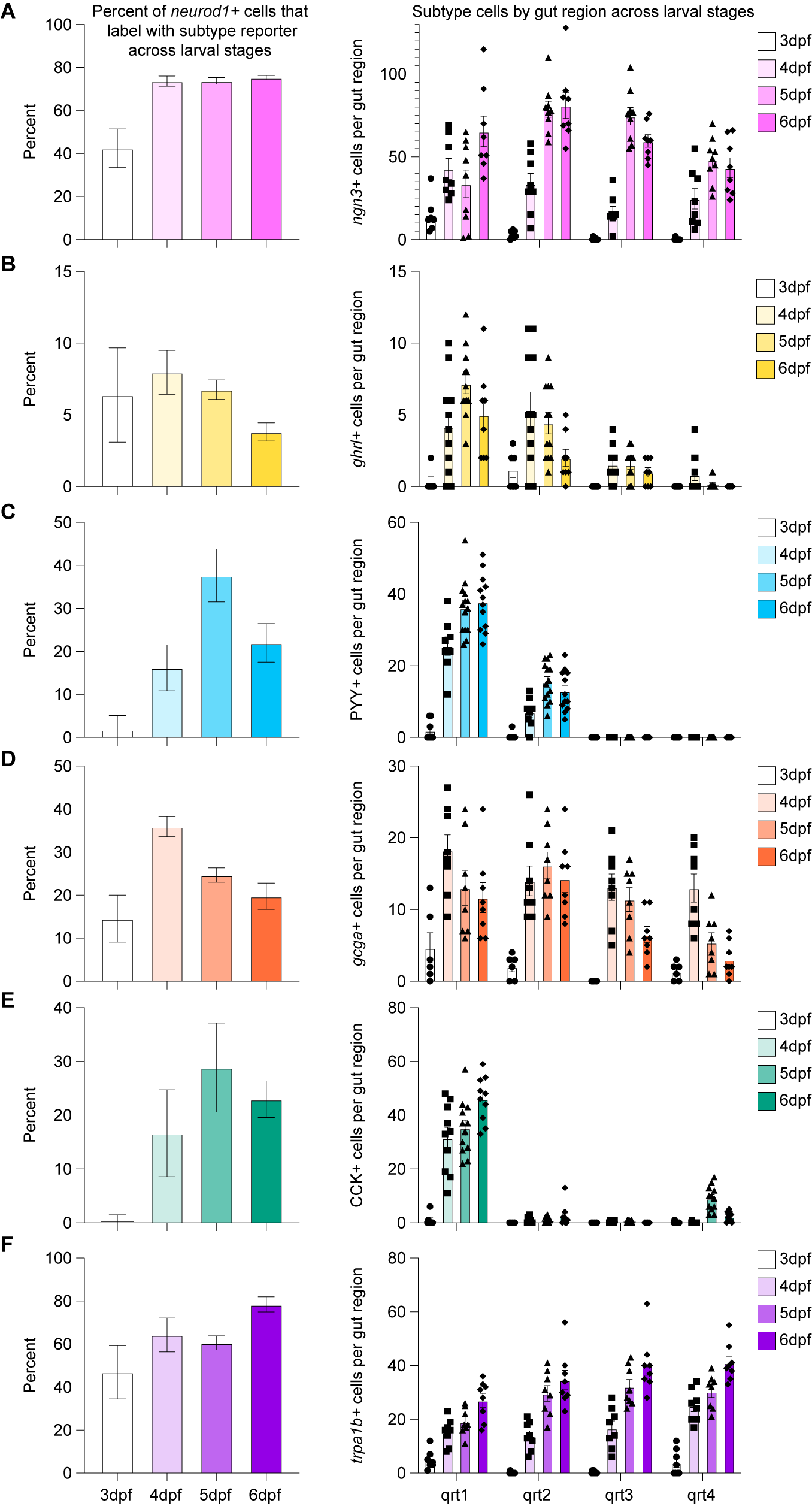

### Sup Fig 8

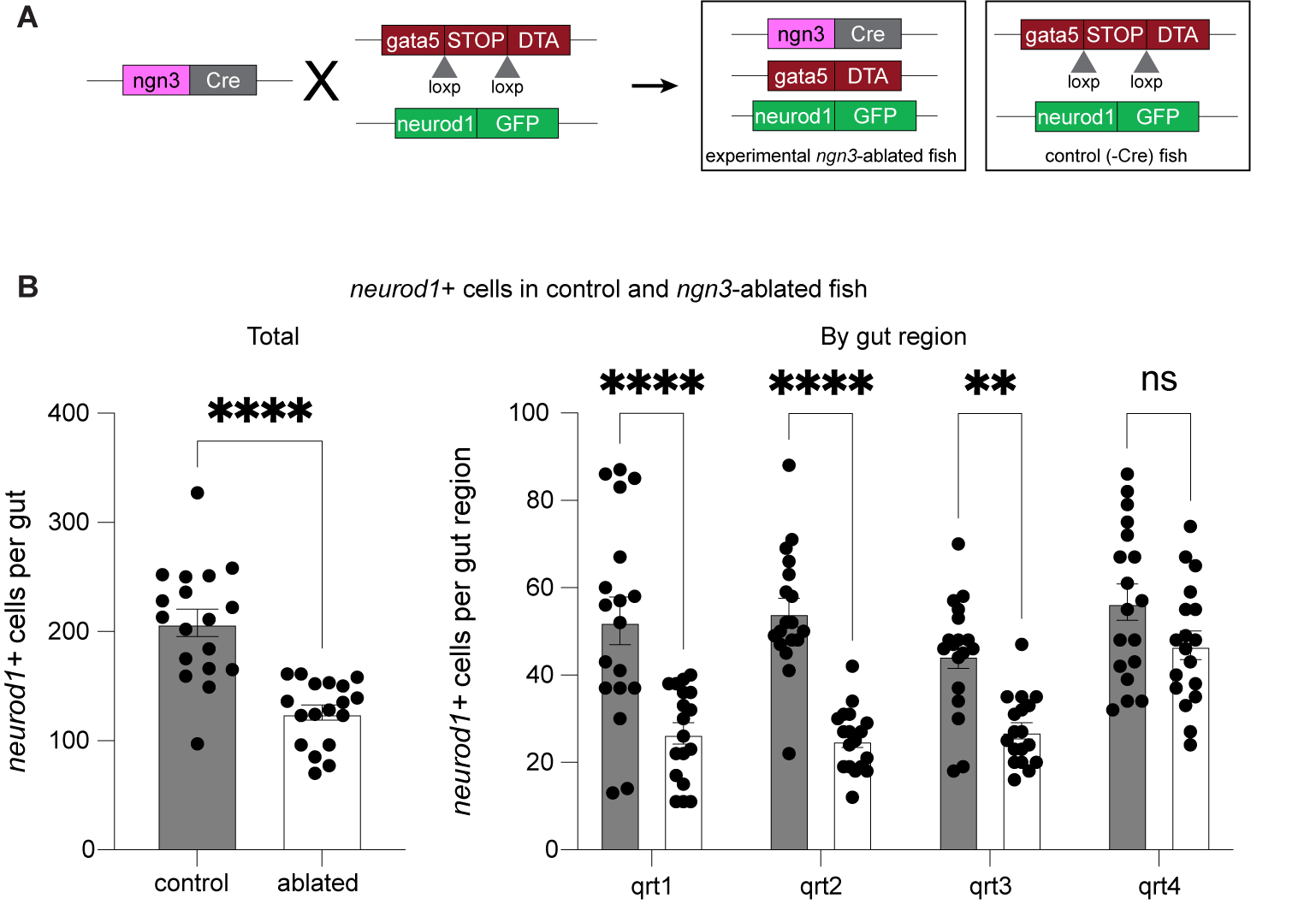

### Sup Fig 9

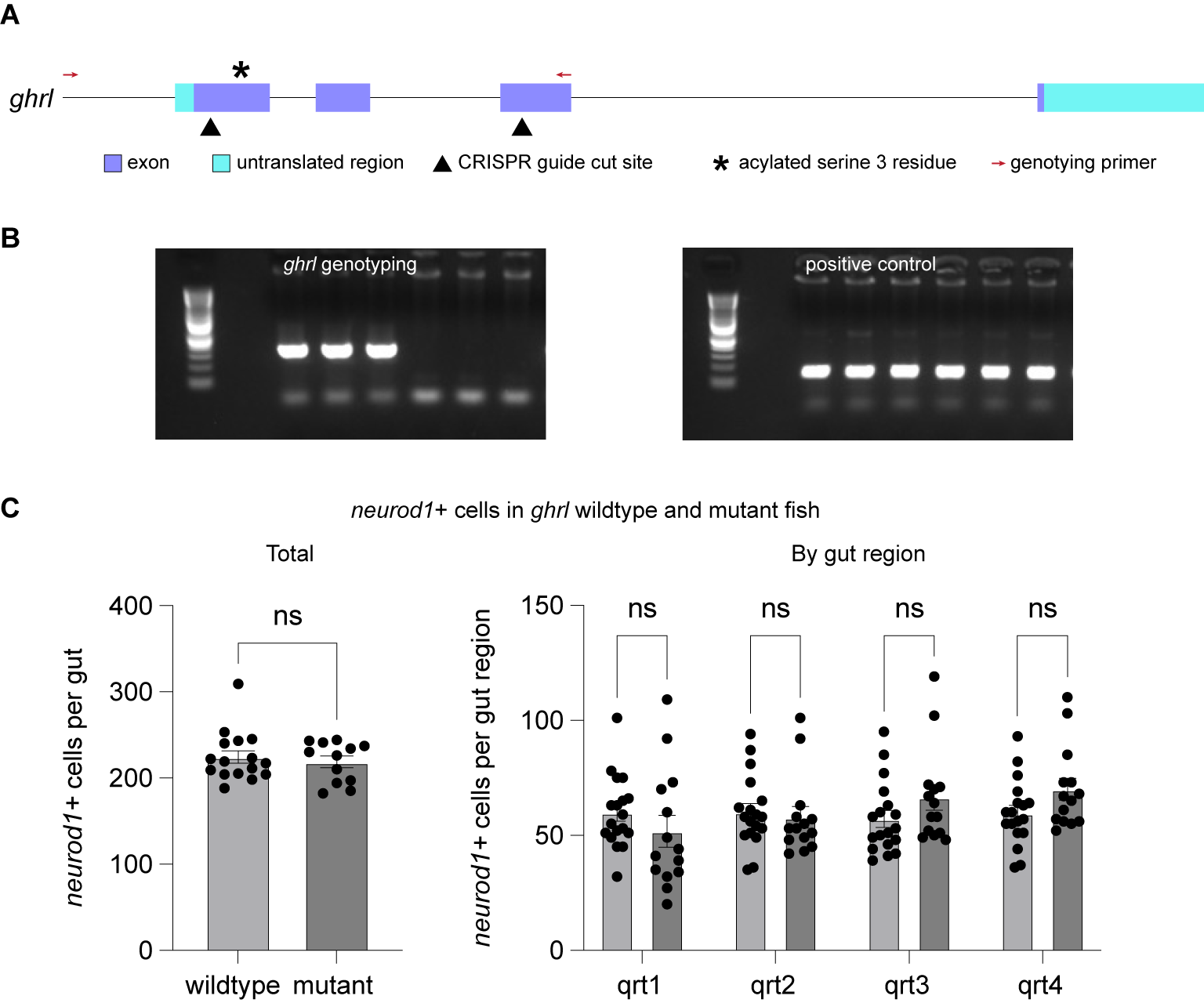

### Sup Fig 10

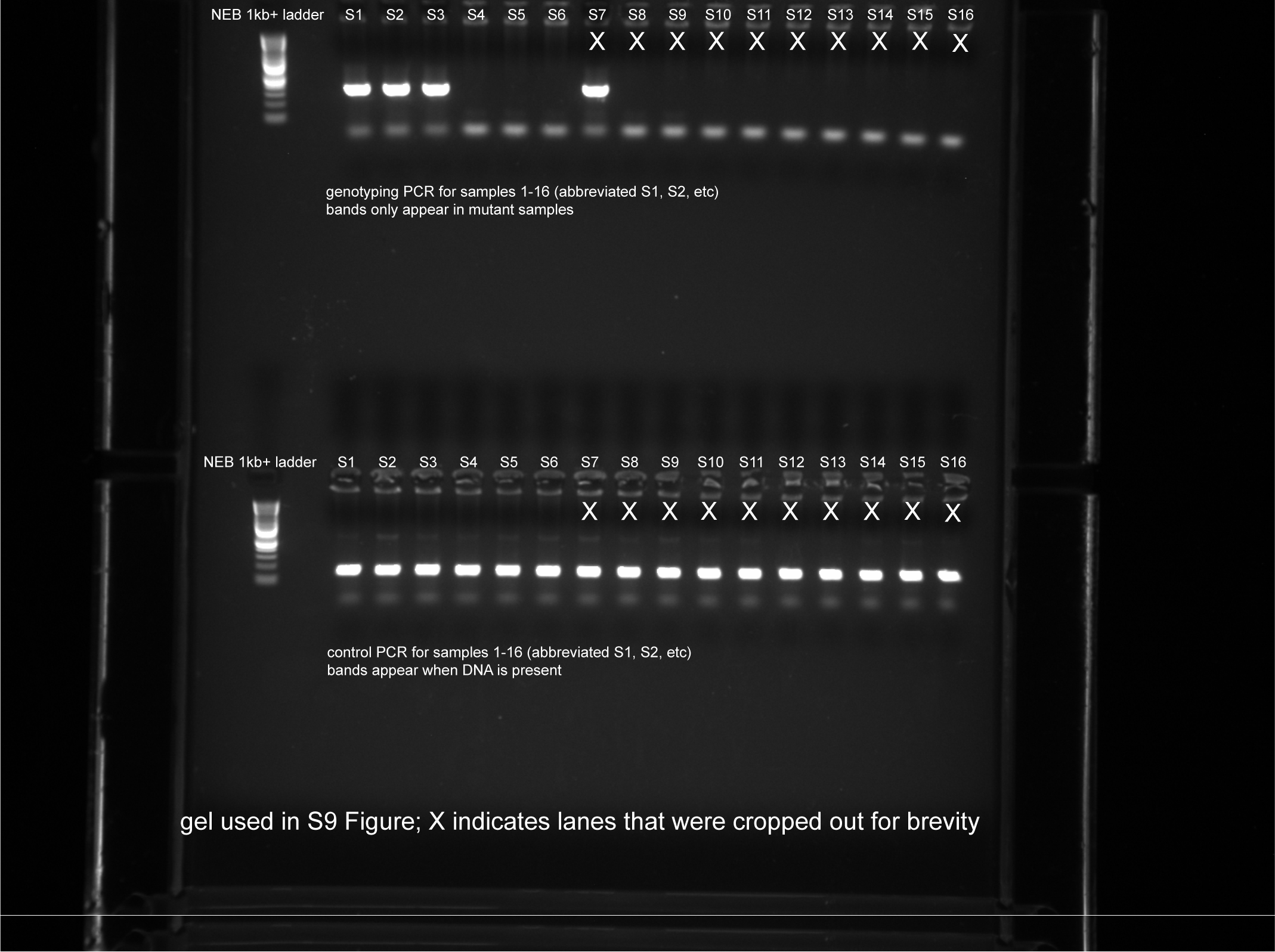
