## Supplementary material for "Intestinal enteroendocrine cell subtype differentiation and hormone production in zebrafish": Sup Fig 3

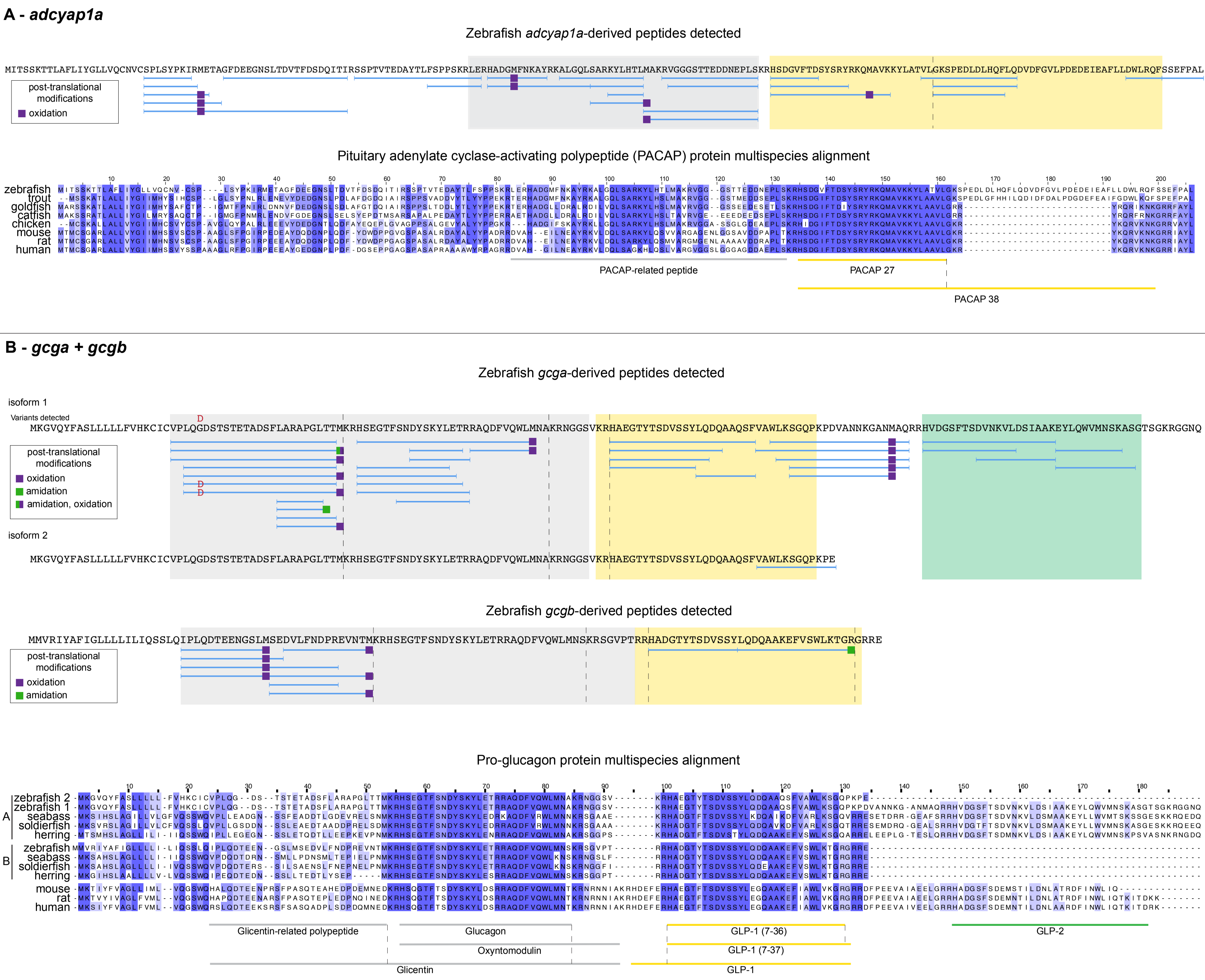

A - adcyap1a
Zebrafish adcyap1a-derived peptides detected
MITSSKTTLAFLIYGLLVQCNVCSPLSYPKIRMETAGFDEEGNSLTDVTFDSDQITIRSSPTVTEDAYTLFSPPSKRLERHADGMFNKAYRKALGQLSARKYLHTLMAKRVGGGSTTEDDNEPLSKRHSDGVFTDSYSRYRKQMAVKKYLATVLGKSPEDLDLHQFLQDVDFGVLPDEDEIEAFLLDWLRQFSSEFPAL
post-translational modifications
oxidation
Pituitary adenylate cyclase-activating polypeptide (PACAP) protein multispecies alignment
zebrafish
trout
goldfish
catfish
chicken
mouse
rat
human
PACAP-related peptide
PACAP 27
PACAP 38
B - gcga + gcgb
Zebrafish gcga-derived peptides detected
isoform 1
Variants detected
MKGVQYFASLLLLLFVHKCICVPLQGDSTSTETADSFLARAPGLTTMKRHSEGTFSNDYSKYLETRRAQDFVQWLMNAKRNGGSVKRHAEGTYTSDVSSYLQDQAAQSFVAWLKSGQPKPDVANNKGANMAQRRHVDGSFTSDVNKVLDSIAAKEYLQWVMNSKASGTSGKRGGNQ
post-translational modifications
oxidation
amidation
amidation, oxidation
isoform 2
MKGVQYFASLLLLLFVHKCICVPLQGDSTSTETADSFLARAPGLTTMKRHSEGTFSNDYSKYLETRRAQDFVQWLMNAKRNGGSVKRHAEGTYTSDVSSYLQDQAAQSFVAWLKSGQPKPE
Zebrafish gcgb-derived peptides detected
MMVRIYAFIGLLLLILIQSSLQIPLQDTEENGSLMSEDVLFNDPREVNTMKRHSEGTFSNDYSKYLETRRAQDFVQWLMNSKRSGVPTRRHADGTYTSDVSSYLQDQAAKEFVSWLKTGRGRRE
post-translational modifications
oxidation
amidation
Pro-glucagon protein multispecies alignment
A zebrafish 2
zebrafish 1
seabass
soldierfish
herring
B zebrafish
seabass
soldierfish
herring
mouse
rat
human
Glicentin-related polypeptide
Glucagon
Oxyntomodulin
Glicentin
GLP-1 (7-36)
GLP-1 (7-37)
GLP-1
GLP-2

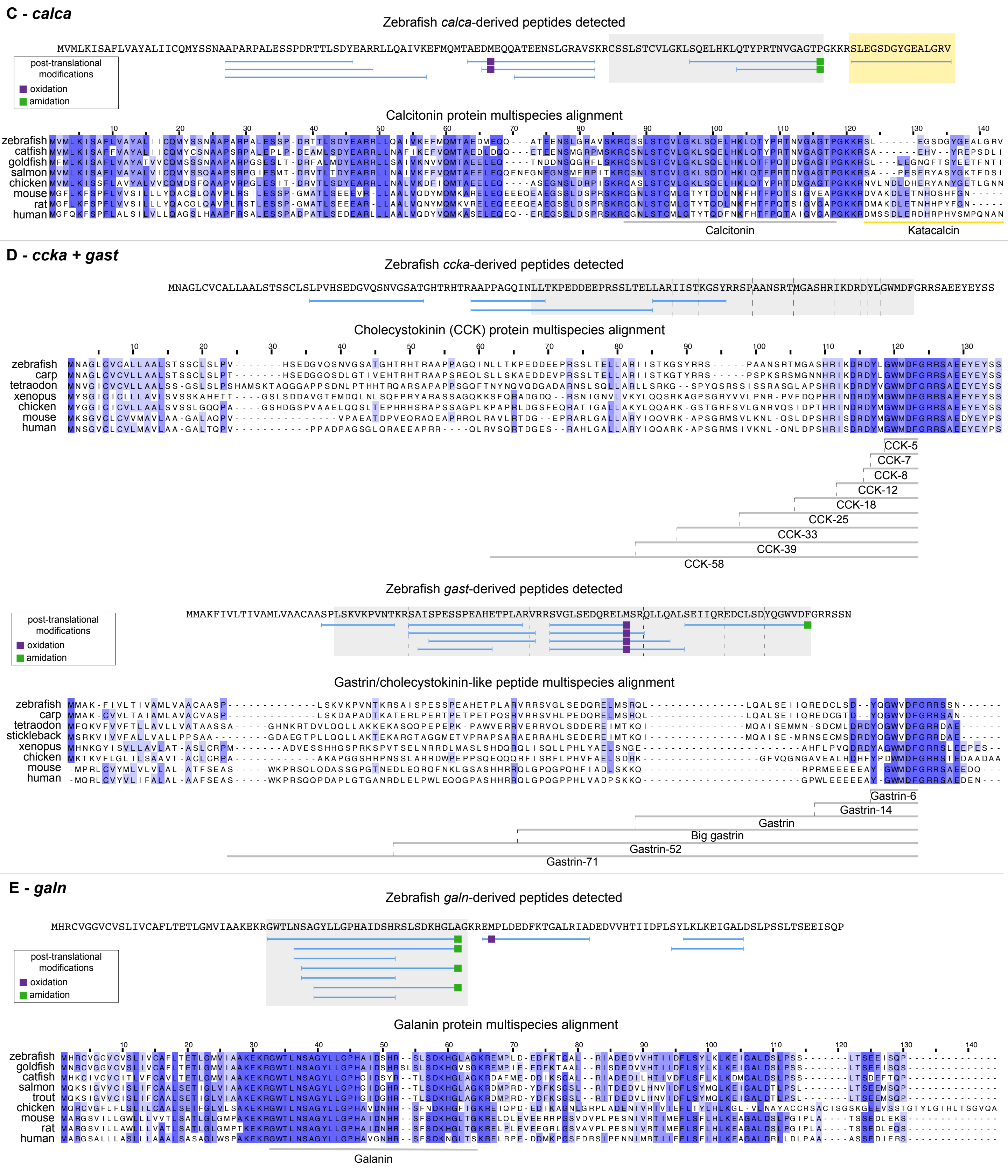
C - calca
Zebrafish calca-derived peptides detected
post-translational modifications
oxidation
amidation
Calcitonin protein multispecies alignment
zebrafish
catfish
goldfish
salmon
chicken
mouse
rat
human
Calcitonin
Katacalcin
D - ccka + gast
Zebrafish ccka-derived peptides detected
Cholecystokinin (CCK) protein multispecies alignment
zebrafish
carp
tetraodon
xenopus
chicken
mouse
human
CCK-5
CCK-7
CCK-8
CCK-12
CCK-18
CCK-25
CCK-33
CCK-39
CCK-58
Zebrafish gast-derived peptides detected
post-translational modifications
oxidation
amidation
Gastrin/cholecystokinin-like peptide multispecies alignment
zebrafish
carp
tetraodon
stickleback
xenopus
chicken
mouse
human
Gastrin-6
Gastrin-14
Gastrin
Big gastrin
Gastrin-52
Gastrin-71
E - galn
Zebrafish galn-derived peptides detected
post-translational modifications
oxidation
amidation
Galanin protein multispecies alignment
zebrafish
goldfish
catfish
salmon
trout
chicken
mouse
rat
human
Galanin

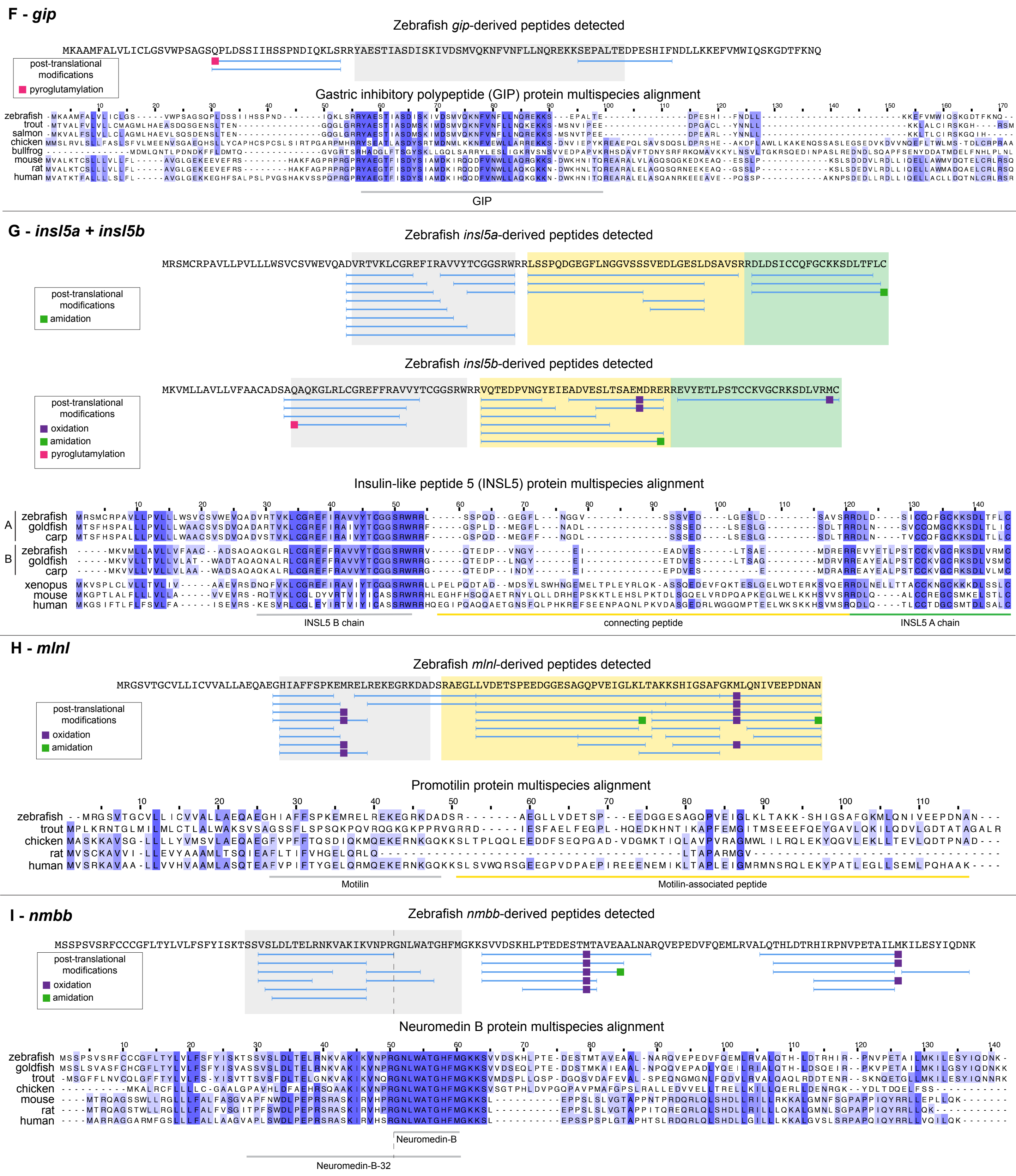

F - gip
Zebrafish gip-derived peptides detected
MKAAMFALVLICLGSVVWPSAGSQPLDSSIIHSSPNDIQKLSRRYAESTIASDISKIVDSMVQKNFVNFLLNQREKKSEPALTEDPESHIFNDLLKKEFVMWIQSKGDTFKNQ
post-translational modifications
pyroglutamylation
Gastric inhibitory polypeptide (GIP) protein multispecies alignment
zebrafish
trout
salmon
chicken
bullfrog
mouse
rat
human
GIP
G - insl5a + insl5b
Zebrafish insl5a-derived peptides detected
MRSMCRPAVLLPVLLLWSVCSVWEVQADVRTVKLCGREFIRAVVYTCGGSRWRRLSSPQDGEGFLNGGVSSSVEDLGESLDSAVSRRDLDSICCQFGCKKSDLTFLC
post-translational modifications
amidation
Zebrafish insl5b-derived peptides detected
MKVMLLAVLLVFAACADSAQAQKGLRLCGREFFRAVVYTCGGSRWRRVQTEDPVNGYEIEADVESLTSAEMDRERREVYETLPSTCCKVGCRKSDLVRMC
post-translational modifications
oxidation
amidation
pyroglutamylation
Insulin-like peptide 5 (INSL5) protein multispecies alignment
A zebrafish
goldfish
carp
B zebrafish
goldfish
carp
xenopus
mouse
human
INSL5 B chain
connecting peptide
INSL5 A chain
H - mlnl
Zebrafish mlnl-derived peptides detected
MRGSVTGCVLLICVVALLAEQAEGHIAFFSPKEMRELREKEGRKDADSRAEGLLVDETSPEEDGGESAGQPVEIGLKLTAKKSHIGSAFGKMLQNIVEEPDNAN
post-translational modifications
oxidation
amidation
Promotilin protein multispecies alignment
zebrafish
trout
chicken
rat
human
Motilin
Motilin-associated peptide
I - nmbb
Zebrafish nmbb-derived peptides detected
MSSPSVSRFCCCGFLTYLVLFSFYISKTSSVSLDLTELRNKVAKIKVNPRGNLWATGHFMGKKSVVDSKHLPTEDESTMTAVEAALNARQVEPEDVFQEMLRVALQTHLDTRHIRPNVPETAILMKILESYIQDNK
post-translational modifications
oxidation
amidation
Neuromedin B protein multispecies alignment
zebrafish
goldfish
trout
chicken
mouse
rat
human
Neuromedin-B
Neuromedin-B-32

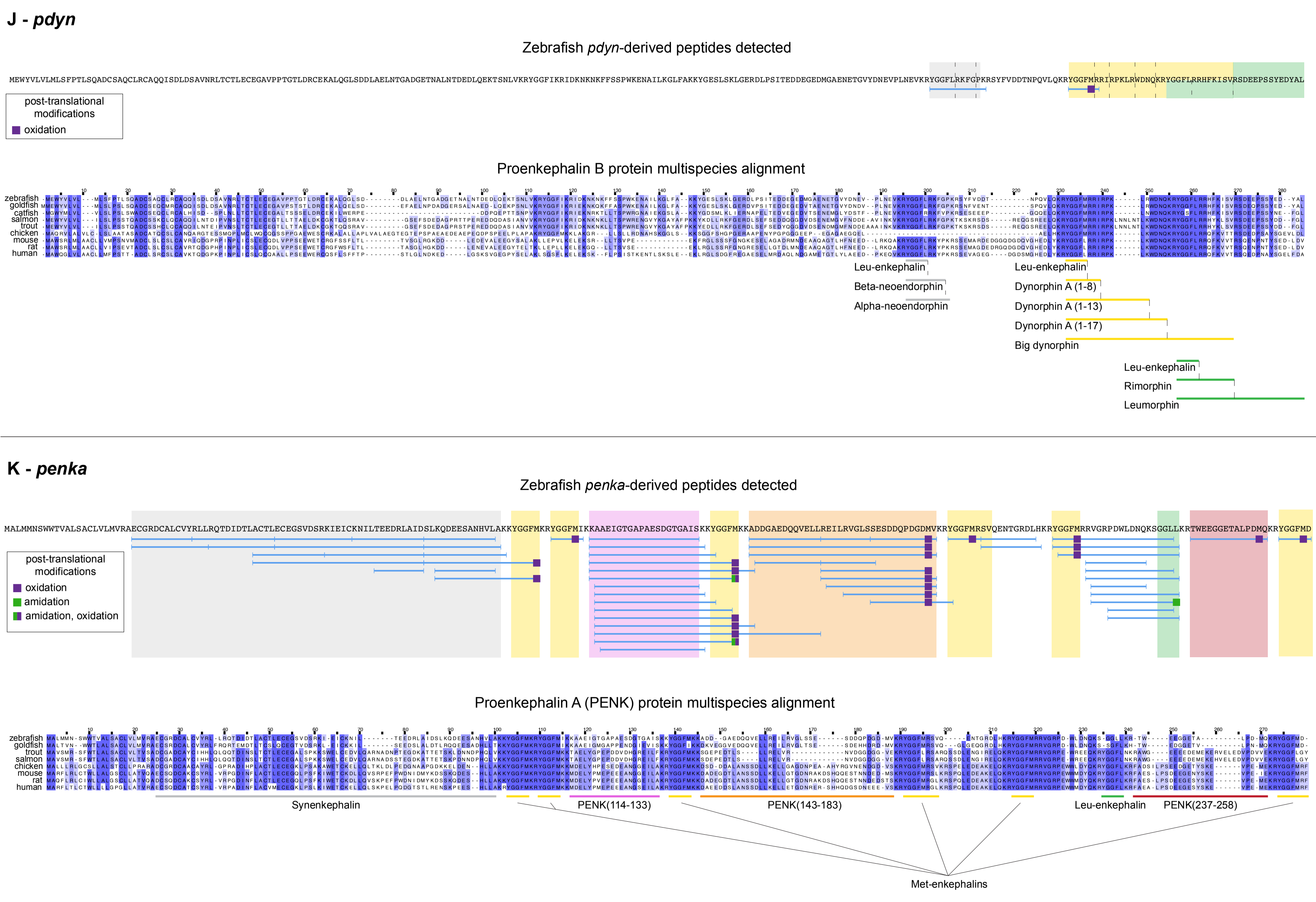
J - pdyn
Zebrafish pdyn-derived peptides detected
MEWYVLVLMLSFPTLSQADCSAQCLRCAQQISDLDSAVNRLTCTLECEGAVPPTGTLDRCEKALQGLSDDLAELNTGADGETNALNTDEDLQEKTSNLVKRYGGFIKRIDKNKNKFFSSPWKENAILKGLFAKKYGESLSKLGERDLPSITEDDEGEDMGAENETGVYDNEVPLNEVKRYGGFLRKFGPKRSYFVDDTNPQVLQKRYGGFMRRIRPKLRWDNQKRYGGFLRRHFKISVRSDEEPSSYEDYAL
post-translational modifications
oxidation
Proenkephalin B protein multispecies alignment
zebrafish
goldfish
catfish
salmon
trout
chicken
mouse
rat
human
Leu-enkephalin
Beta-neoendorphin
Alpha-neoendorphin
Leu-enkephalin
Dynorphin A (1-8)
Dynorphin A (1-13)
Dynorphin A (1-17)
Big dynorphin
Leu-enkephalin
Rimorphin
Leumorphin
K - penka
Zebrafish penka-derived peptides detected
MALMMNSWWTVALSACLVLMVRAECGRDCALCVYRLLRQTDIDTLACTLECEGSVDSRKIEICKNILTEEDRLAIDSLKQDEESANHVLAKKYGGFMKRYGGFMIKKAAEIGTGAPAESDGTGAISKKYGGFMKKADDGAEDQQVELLREILRVGLSSESDDQPDGDMVKRYGGFMRSVQENTGRDLHKRYGGFMRRVGRPDWLDNQKSGGLLKRTWEEGGETALPDMQKRYGGFMD
post-translational modifications
oxidation
amidation
amidation, oxidation
Proenkephalin A (PENK) protein multispecies alignment
zebrafish
goldfish
trout
salmon
chicken
mouse
rat
human
Synenkephalin
PENK(114-133)
PENK(143-183)
Leu-enkephalin
PENK(237-258)
Met-enkephalins

**L - *pyyb***

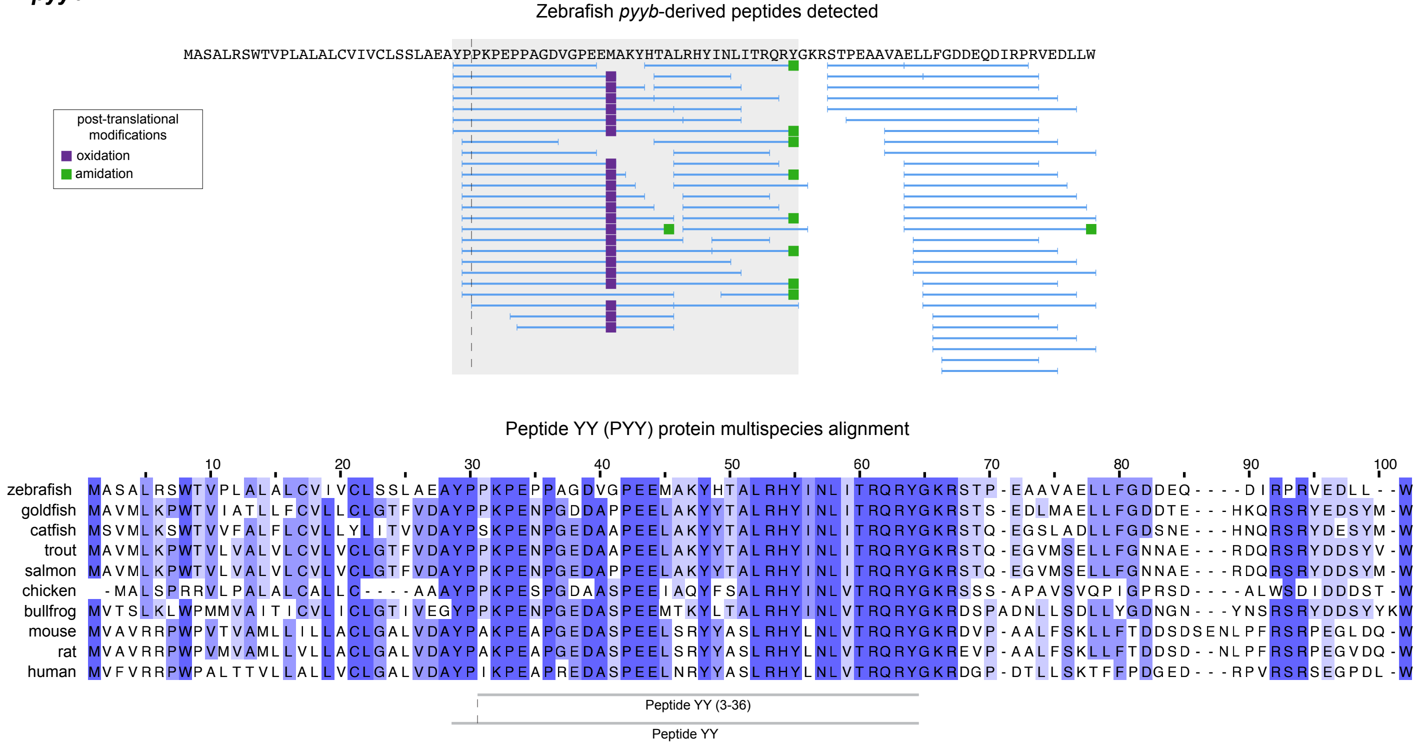


**M - *sst2* + *sst1.2***

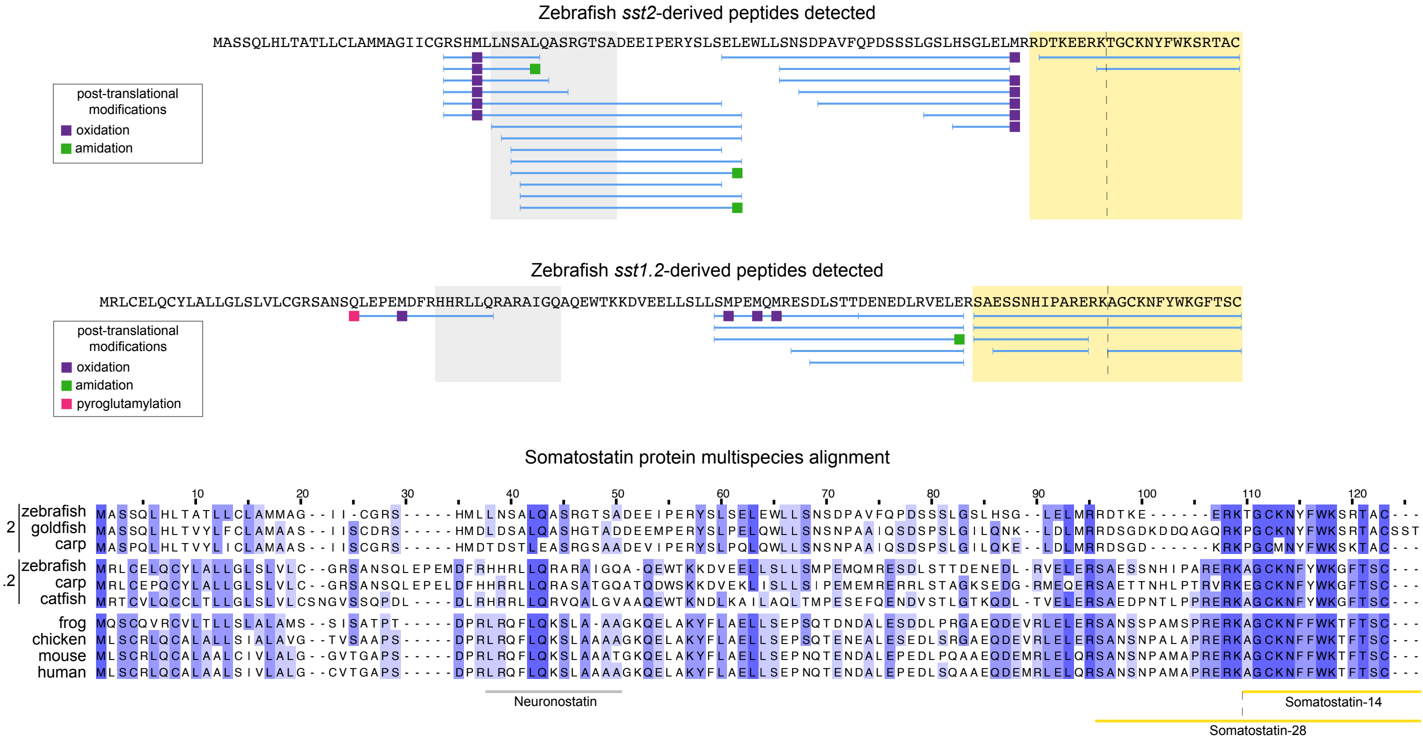

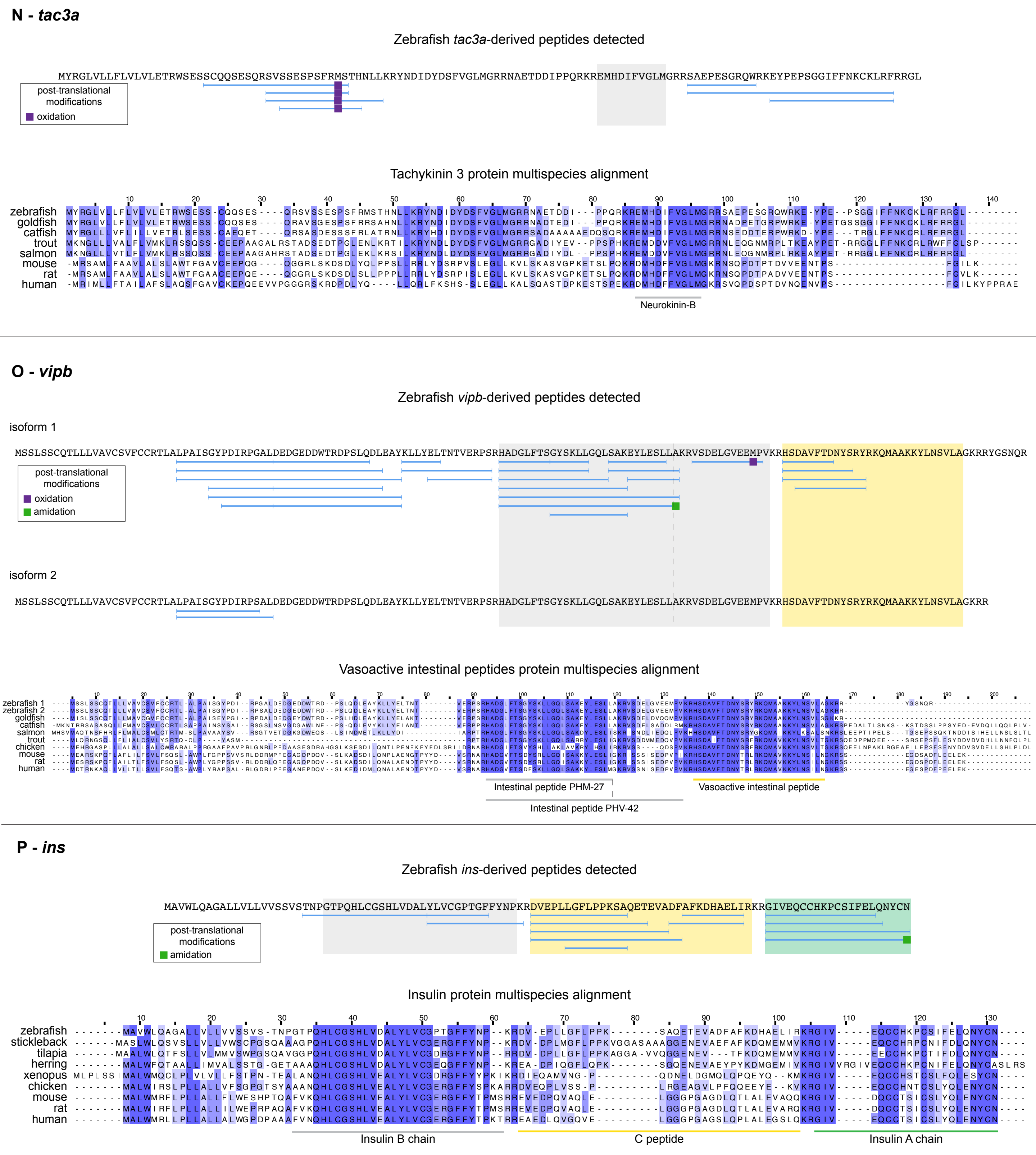
N - tac3a
Zebrafish tac3a-derived peptides detected
MYRGLVLLFLVLVLETRWSESSCQQSESQRSVSSESPSFRMSTHNLLKRYNDIDYDSFVGLMGRRNAETDDIPPQRKREMHDIFVGLMGRRSAEPESGRQWRKEYPEPSGGIFFNKCKLRFRRGL
post-translational modifications
oxidation
Tachykinin 3 protein multispecies alignment
zebrafish
goldfish
catfish
trout
salmon
mouse
rat
human
Neurokinin-B
O - vipb
Zebrafish vipb-derived peptides detected
isoform 1
MSSLSSCQTLLLVAVCSVFCCRTLALPAISGYPDIRPGALDEDGEDDWTRDPSLQDLEAYKLLYELTNTVERPSRHADGLFTSGYSKLLGQLSAKEYLESLLAKRVSDELGVEEMPVKRHSDAVFTDNYSRYRKQMAAKKYLNSVLAGKRRYGSNQR
post-translational modifications
oxidation
amidation
isoform 2
MSSLSSCQTLLLVAVCSVFCCRTLALPAISGYPDIRPSALDEDGEDDWTRDPSLQDLEAYKLLYELTNTVERPSRHADGLFTSGYSKLLGQLSAKEYLESLLAKRVSDELGVEEMPVKRHSDAVFTDNYSRYRKQMAAKKYLNSVLAGKRR
Vasoactive intestinal peptides protein multispecies alignment
zebrafish 1
zebrafish 2
goldfish
catfish
salmon
trout
chicken
mouse
rat
human
Intestinal peptide PHM-27
Vasoactive intestinal peptide
Intestinal peptide PHV-42
P - ins
Zebrafish ins-derived peptides detected
MAVWLQAGALLVLLVVSSVSTNPGTPQHLCGSHLVDALYLVCGPTGFFYNPKRDVEPLLGFLPPKSAQETEVADFAFKDHAELIRKRGIVEQCCHKPCSIFELQNYCN
post-translational modifications
amidation
Insulin protein multispecies alignment
zebrafish
stickleback
tilapia
herring
xenopus
chicken
mouse
rat
human
Insulin B chain
C peptide
Insulin A chain

**Q - *pcsk1nl***

Zebrafish *pcsk1nl*-derived peptides detected

MSGLLSSSLLFFLSLTLFQIHTPEAKPLSAMRGGVRSYDVPVRLRRELRDSVPYEAQMISYPSADFKSRSNDYYPSEVLRAQGLGQALQRLVESDQRREQEAAYLASMLRLLNEAQGTGQGRAGDQEEEGGDFQGPYPPDYDETEQA

VSMAKPQASWQGLLDPQLTQALLNRYKQERLVQAGLAPAANRIPEREQDKDQEMLRYLVEKILSSLASGGNQGSSSNPRAKRDLSAVSSIERPIKPALKRSRRSLDSAPGPQAEASLLRVKRLDDDDDVDEDAVAGQSNGTPHIGLQ

RMKRIDTDLPPPKKHSRKRRALSYDPALIAQHILQYLPA

---

**R - *pcsk1***

Zebrafish *pcsk1*-derived peptides detected

MEERCRPMLLCSVLALVCALVSSAVDRQYLNEWAVEIPGGVQNARSIADEFGYQLVRQIGALENHYLFKRHSHPSRTKRSADHITKRLSEDDRVSWAEQQYEKRRAKRAPLGVECKDCSVDKLFDDPMWNQQWYLQDTRTSSSLPKL

DLHVIPVWKKGITGKGVVITVLDDGLEWNHTDIYPNYDPAASYDFNDNDPDPFPRYDSTNENKHGTRCAGEIAMQADNNKCGVGVAYNSKVGGIRMLDGIVTDAIEASSIGYNPDHVDIYSASWGPNDDGKTVEGPGRLAQKAFEYG

IQKGRGGKGSIFVWASGNGGRQGDNCDCDGYTDSLYTISISSASQQGLSPWYAEKCSSTLATAYSSGDYTDQRITSADLHNECTETHTGTSASAPLAAGIFALALEQNPDLTWRDLQHLVVWTSEFDPLANNPGWKRNGAGLMVNSR

FGFGLLNAKALVDLADPKVWKHVPEKKQCIVRDETFQPRPLKAAGEISIEIPTKACAGQANSVMSLEHVQVEVSIEYTRRGDLHITLTSPSGTTTVLLAERERDTSSNGFRNWAFMSVHTWGENPTGTWILKITDTSGRMENEGQII

SWKLILHGTSEKPEHMKKARVYTPYNAVQNDRRGVEPMEDMMKQEAPTPALKPEIPNQAIPDSPPSTASMSIPEVNSGPSTANLALLRLLQSAFNRQMPAAPPKSFQQERIPPQKLYQALDLLNRYRASQNGLFNDYSDNFYRAQPY

RHRDDRLLQALFDMLRDDQQ

---

**S - *pcsk2***

Zebrafish *pcsk2*-derived peptides detected

MRRFRGHHRTAPALFLTLIGALMMSAAAEDELFITGHFLVQMREAAPEDAQKLALEHGFESARKLPFGEDLFHFYPLEMPKTRRKRSLHQQHHRLASDHRVKNVFPQEGFGRQKRGYRDLPNTDVNMSDPLFTKQWYLINTGQADGT

PGLDLNVAEAWSLGFTGKGVTIAIMDDGIDYLHPDLASNYNAEASFDFSSNDPYPYPRYTDDWFNSHGTRCAGEVSAVSNNNICGVGVAYNSKVAGIRMLDQPFMTDIIEASSISHMPQVIDIYSASWGPTDDGKTVDGPRELTLQA

MADGVNKGRGGKGSIYVWASGDGGSYDDCNCDGYASSMWTISINSAINDGRTALYDESCSSTLASTFSNGRKRNPEAGVATTDLYGNCTLRHSGTSAAAPEAAGVFALALEANPNLTWRDLQHLTVLTSKRNKLHDEVHQWRRNGVG

LEFNHLFGYGVLDAGGMVKLARDWKTVPERFHCVAGSMQDIHKIQSGNKLLLSISTDACQGKDNFVRYLEHVQAVITVNASRRGDLNINMTSPMGTKSILLSRRPRDDDAKVGFDKWPFMTTHTWGEDPRGPWLLEVGFQSQSDMQS

GLLKEWTLMLHGTQSAPYIDQVVRDYQSKLAMSKKEELEEELDEAVERSLKSLLSKNN

### T - *scg2a*

Zebrafish *scg2a*-derived peptides detected

MSSSSSSSRCCAAGVFLVPSLLLLPLLLSLIHTAHGATLREHRLSGTEPVSYGPPSQLRPPPSAEMLRALRYIQSLSERTPADPDQSDQDAEDDMESVRSVLKMASPTRREEKDNTQELLQAVLTTLQQTEEHMKTPQKVIAAPRYH

QFARPKQQVKVDTDDGNYGRNSWAENRRRYREYPIMFEDDQPLKRTNENAEEQYTPQKLATLQSVFEELSGIASSKTNTKRSDDDEDDEDLYRQRKMVLEDIMGTDDWTPLEEQMESEEEERERHGFTRNLEDDDEDDDGKRSLQSD

WLQTQREEEEPEDMAKLVDYYLLQMLEKKEQEQQKRQEEDEEEEVEKKDEEEENVEEREVKPMQSLSEILKISQKLRIPPEELLQLLRNENRKDLPGRTGYAHKTFPSAPHRRPIETSDIAQDILSILELASAAQQNQRPTQARNER

YYERQTSRDDYDDTAGEEDELANYLATEMQRHTRMAPLRDEDQPVNLQNYYEKPPAEHENAATGIDNSTMMKILRLLDPESDDADETDSDAGEKVPEM

### U - *scg2b*

Zebrafish *scg2b*-derived peptides detected

MMLSLPKLSAGGVVVLLATLLHTLTVQGASVRHHRLRGGDQGGFLAPSSDMIKALEYIESLKQRADGPESPTGDYDEVDKFRFLVQLASLQDENTPTHEDATRWPDNKVPQWVRSLLRVLDQAGESPESQAAGNERRLHKTRRPVAD

GESPAGDYAGFVKPHKKYPLMFEDEENGRDNKRATEDLDEQYTPQSLANMRSIFEELGKLSAAQKRDDEEDGEDDDDLYRVRNAAYEDVTGGEEWVPLEEQLETEELVKGSHEEYKRALGDISEQGMENMERRGEEEDENPDDDTKL

VDYYLLKVLEMTDQAQKRDLMEGRRRLLSRPSLIDPRAIKQLLSAISMKLQVPPEDLVGMLFMEETRKQQQRLPEPQLARNPSQPRYKSRVIKYYNGRQPEVTVSDIPHDVKTEDILKVLGLGNLANKNAKFSLLKQRPYKTAMTNY

FNPNGRRGSLFLSELNKAPSKRKDDYDDDDAVDEDEESTFLAAKLLTEYPDTSSSNRKRAIDSAANGQLPYELYEEAMKDFFDQVDNGKSALAKRDTQGKEEPEGPQKPPAQDPAQETVDQTPPESGTEDGKEYHGKIVAGM

### V - *scg3*

Zebrafish *scg3*-derived peptides detected

Variants detected

E A

MASKRLGFVVVLALVCQHINAFPTPAGPDDKYNRELTEEKPLEQQIAEADSIRKTESKPTPPAEEETNSEDDDITFLKALAEKSKESNNETPISDSANERLGADDTDSTKNRRLADDYDSTKNGMDYKYQDDPESFRQLDGTPLTAQ

E

A

A

post-translational modifications

oxidation

amidation

DIVQKIANKIYEEDDRGVFDRIVSKLLKLGLITDSQAETLEYEVAEALQDLITKNAKDNMIGDRSVDYPVSAAKDTQDEEDRPSPRFDEEDEVENEGGDDANGDEPQEEESRDDTVRSDDSWDAVSDGNDRNELNPEDGLQDLQFFP

NFYRLLKSLDSEQDKEERETLITIMKTLIDFVKMMVKYGTITPEEGVTYLENLDAMIALQTKNKLGKSLVPLSITPPTGKAADDDDNTKTEAAKMQKEYESLKDSTKDVQTAAEISHPGKSESYLEAIRKNIEWLKKHNKEGNKEDY

DLSKLRDFMDQQVDSYIDKGILEKDEGDVIKRIYGSL

### W - *scg5*

Zebrafish *scg5*-derived peptides detected

MGSTSKMTHTVFLSVIFSLIAVVSGRSPRTVDQVSEADIQRLLHGVMEQLGIARPRVEYPAHQATNIVGPLSIQGGAHEGLQHLGPYGNIPNIVAELTGDSVPKDFSEDHGYPDPPNPCPLGKTAADGCLENSPDTAEFSREF

post-translational modifications

oxidation

QKHQHLFDPEHDYPALAKWNKGLLYEKLKGGPKRRKRSVPNPYLMGQRLDNVVAKKSVPHFPEEDEVETATSNIKT

### X - *scgn*

Zebrafish *scgn*-derived peptides detected

MDSAFANLDAAGFLQIWQHFDADDNGYIEGKELDDFFRHMLKKLQPKDKITDERVQQIKKSFMSAYDATFDGRLQIEELANMILPQEENFLLIFRREAPLDNSVEFMKIWRKYDADSSGYISAAELKNFLKDLFLQHKKKIP

post-translational modifications

oxidation

PNKLDEYTDAMMKIFDKNKDGRLDLNDLARILALQENFLLQFKMDASSQVERKRDFEKIFAHYDVSRTGALEGPEVDGFVKDMMELVRPSISGGDLDKFRECLLTHCDMNKDGKIQKSELALCLGLKHKP
