## Supplementary material for "Intestinal enteroendocrine cell subtype differentiation and hormone production in zebrafish": Sup Fig legends

**Supplemental information**

**S1 Table. EEC scRNA-seq markers.**

**S2 Table. Zebrafish EEC peptidomics.**

**S1 Fig. Creation of joint larval and adult intestinal secretory cell scRNA-seq dataset.**

**(A)** Raw scRNA-seq data of the adult zebrafish intestine from [49] was clustered and evaluated for expression of secretory cell markers. Clusters showing strong expression of these markers and intestinal epithelial markers but minimal pancreatic endocrine cell markers were selected for subsequent analysis and are outlined with red boxes. **(B)** We similarly processed scRNA-seq data of the larval zebrafish intestine from [48] and selected clusters outlined in red for subsequent analysis. **(C)** UMAP of the joint adult and larval dataset generated by integrating and re-clustering cells identified in panels A and B. Clusters circled with a dashed line were identified as EECs and secretory progenitors based on expression shown in **(D)** and were selected for subsequent integration and re-clustering to form the dataset shown in Fig 1.

**S2 Fig. Additional characterization of joint adult and larval secretory progenitor and EEC scRNA-seq dataset.**

**(A)** Expression Dotplot showing secretory progenitor markers are enriched in clusters 7-10. **(B)** UMAP colored by dataset of origin for each cell with adult cells in green and larval cells in blue. **(C)** Stacked bar plot showing the percentage of cells in each EEC cluster from the adult, larval, or combined dataset. **(D)** Heatmap showing the z-scored expression of the 10 most highly enriched markers for each EEC cluster. **(E)** Heatmap showing the z-scored expression of genes annotated as receptors by [168] that were significantly enriched in both larval and adult cells in each cluster. **(F)** Heatmap showing the z-scored expression of genes annotated as ligands by [168] that were significantly enriched in both larval and adult cells in each cluster. **(G)** Table of major differences between the samples that generated the larval and adult scRNA-seq datasets.

**S3 Fig. Zebrafish EEC peptide atlas.**

In all panels, the primary amino acid sequence of the gene of interest is shown at the top in black with any identified missense variants indicated in red above. Blue horizontal lines below the amino acid sequence represent the individual peptides detected in our study with small vertical lines denoting the stop and start of each peptide. Different colored squares represent various posttranslational modification detected. Shading labels regions aligning to Uniprot-annotated peptides in humans. In cases where multiple peptides are known to be processed from the same sequence, dashed lines indicate different cleavage sites. Below the detected peptides is a multispecies protein alignment where amino acids are color coded based on their percent identity match across all the reported species with darker coloring indicating a more conserved residue. Human processed peptide annotations taken from Uniprot are labeled below with horizontal lines. The color of these lines corresponds to the color of the shading of the aligned zebrafish amino acids above. Note that many of the peptide cleavage sites occur at dibasic residues (i.e., RR/KR/KK) consistent with cleavage by prohormone convertase enzymes [120]. More detailed information about the peptides shown in this figure are available in S2 Table. **(A)** Zebrafish *adcyap1a-*derived peptides detected in EECs. Alignment of the primary amino acid sequence of Pituitary adenylate cyclase-activating polypeptide protein includes zebrafish (Uniprot Q98TU3), trout (Uniprot A0A8C7QBD4), goldfish (Uniprot A0A6P6RC36), catfish (Uniprot Q90XZ4), chicken (Uniprot Q58FG9), mouse (Uniprot O70176), rat (Uniprot A6KFB2), human (Uniprot P18509). **(B)** Zebrafish *gcga-* and *gcgb-*derived peptides detected in EECs. As *gcga* and *gcgb* are both orthologs of human *GCG*, they are reported together. Peptides aligning to multiple isoforms of *gcga* were detected. Alignment of the primary amino acid sequence of Pro-glucagon protein includes zebrafish *gcga* isoform 2 (Uniprot F1RD10), zebrafish *gcga* isoform 1 (Uniprot A0A0R4IS85), seabass *gcga* (Uniprot A0A8C4IA77), soldierfish *gcga* (Uniprot A0A667XLC6), herring *gcga* (Uniprot A0A6P3WDK7), zebrafish *gcgb* (Uniprot B0R1C3), seabass *gcgb* (Uniprot A0A8P4K489), soldierfish *gcgb* (Uniprot A0A668AYA2), herring (Uniprot A0A6P3W7W6), mouse *Gcg* (A2AS86), rat *Gcg* (Uniprot A6HLV7), human (Uniprot P01275). Of note, the arginine 36 residue in GLP-1 is commonly amidated [116,117], as seen in the *gcgb*-derived peptide reported here. **(C)** Zebrafish *calca*-derived peptides detected in EECs. Alignment of the primary amino acid sequence of Calcitonin protein includes zebrafish (Uniprot F1QIK6), catfish (Uniprot A0A2D0RV9), goldfish (Uniprot A0A6P6MFE7), salmon (Uniprot A0A1S3KMJ8), chicken (Uniprot P07660), mouse (Uniprot P70160), rat (Uniprot P01257), and human (Uniprot P01258). **(D)** Zebrafish *ccka-* and *gast-*derived peptides detected in EECs. As Cholecystokinin (CCK) and Gastrin are closely related, they were aligned together, but are displayed separately for ease of visualization. Of note, the zebrafish *gastrin* gene is currently named as *CR556712.1* or **LOC100536965**, but due to evidence from published phylogenetic studies [161–163] and protein sequence alignment, we refer to it as *gastrin* (see Methods)*.* Alignment of the primary amino acid sequence of CCK and Gastrin includes zebrafish *ccka* (Uniprot E9QEB3), carp *ccka* (Uniprot A0A9R1SKH2), tetraodon *ccka* (Uniprot Q8AXP6), xenopus *cck* (Uniprot A0A803J912), chicken *cck* (Uniprot Q9PU41), mouse *Cck* (Uniprot P09240), human *CCK* (Uniprot P06307), zebrafish *gast* (Uniprot A0A8M6Z182), carp *gast* (Uniprot A0A8C1B8C9), tetraodon *gast* (Uniprot Q8AXP5), stickleback *gast* (Ensembl ENSGACT00000024275.2), xenopus *gast* (Uniprot F6W277), chicken (Ensembl ENSGALT00000043859.2), mouse *Gast* (Uniprot P48757), human *GAST* (Uniprot P01350). **(E)** Zebrafish *galn*-derived peptides detected in EECs. Alignment of the primary amino acid sequence of Galanin includes zebrafish (Uniprot E7EZ53), goldfish (Uniprot Q7ZT91), catfish (Uniprot A0A2D0RH49), salmon (Uniprot A0A1S3KNE4), trout (Uniprot A0A060WZ82), chicken (Uniprot A0A8V0ZZ93), mouse (Uniprot P47212), rat (Uniprot A6HYK9), human (Uniprot P22466). **(F)** Zebrafish *gip*-derived peptides detected in EECs. Alignment of the primary amino acid sequence of Gastric inhibitory polypeptide includes zebrafish (Uniprot A1DPK4), trout (Uniprot A0A060W6J5), salmon (Uniprot A0A1S3NHD2), chicken (Uniprot A0A8V1AAS2), bullfrog (Uniprot A0A2G9RZ64), mouse (Uniprot P48756), rat (Uniprot A6HIC4), human (Uniprot P09681). **(G)** Zebrafish *insl5a-* and *insl5b-*derived peptides detected in EECs. As *insl5a* and *insl5b* are both orthologs of human *INSL5*, they are reported together. Alignment of the primary amino acid sequence of Insulin-like peptide 5 includes zebrafish *insl5a* (Uniprot Q2VT44), goldfish *insl5a* (Uniprot 0A6P6KC48), carp *insl5a* (Uniprot A0A8C1HD46), zebrafish *insl5b* (Uniprot A0ZYT5), goldfish *insl5b* (Uniprot A0A6P6JU13), carp *insl5b* (Uniprot A0A9J8CMM8), xenopus *insl5* (Uniprot A0A6I8SQK1), mouse *Insl5* (Uniprot Q9WUG6), human *INSL5* (Uniprot Q9Y5Q6). **(H)** Zebrafish *mlnl*-derived peptides detected in EECs. Alignment of the primary amino acid sequence of Promotilin includes zebrafish (Uniprot E9QFU1), trout (Uniprot A0A060WHA4), chicken (Uniprot A0A8V1AED3), rat (Uniprot A8IRI0), human (Uniprot P12872). **(I)** Zebrafish *nmbb*-derived peptides detected in EECs. Alignment of the primary amino acid sequence of Neuromedin B includes zebrafish (Uniprot B3DFU2), goldfish (Uniprot A0A6P6PMY7), trout (Uniprot A0A8K9XPV7), chicken (Uniprot A0A8V0ZQC9), mouse (Uniprot Q9CR53), rat (Uniprot A6JCD9), human (Uniprot P08949). **(J)** Zebrafish *pdyn*-derived peptides detected in EECs. Alignment of the primary amino acid sequence of Proenkephalin B includes zebrafish (Uniprot Q6JT77), goldfish (Uniprot A0A6P6NSK2), catfish (Uniprot E3TFZ0), salmon (Uniprot B5X928), trout (Uniprot A0A060WJX8), chicken (Uniprot A0A8E7KML8), mouse (Uniprot O35852), rat (Uniprot F1M7S3), human (Uniprot P01213). **(K)** Zebrafish *penka*-derived peptides detected in EECs. Alignment of the primary amino acid sequence of Proenkephalin A includes zebrafish (Uniprot A8E7S2), goldfish (Uniprot A0A6P6MMG9), trout (Uniprot A0A8C7NZE4), salmon (Uniprot B5X739), chicken (Uniprot E1C652), mouse (Uniprot P22005), rat (Uniprot A6JFM8), human (Uniprot P01210). **(L)** Zebrafish *pyyb*-derived peptides detected in EECs. Alignment of the primary amino acid sequence of Peptide YY (PYY) includes zebrafish (Uniprot E7F0L6), goldfish (Uniprot A0A6P6PFJ3), catfish (Uniprot W5UQI6), trout (Uniprot A0A060XNM3), salmon (Uniprot A0A1S3S6S3), chicken (Uniprot A0A0S3UPI6), bullfrog (Uniprot A0A2G9P7P3), mouse (Uniprot H3BK86), rat (Uniprot F1LSR6), human (Uniprot P10082). **(M)** Zebrafish *sst2-* and *sst1.2*-derived peptides detected in EECs. As both *sst2* and *sst1.2* are orthologs of human *SST*, they are reported together. *sst2* is considered to be part of SST family 4 and *sst1.2* is considered to be part of SST family 3 [169]. Alignment of the primary amino acid sequence of Somatostatin includes zebrafish *sst2* (Uniprot Q9DDE4), goldfish *sst2* (Uniprot A0A6P6P309), carp *sst2* (Uniprot A0A8C1E251), zebrafish *sst1.2* (Uniprot E7FFY9), carp *sst1.2* (Uniprot A0A8C1BWP9), catfish *sst1.2* (Uniprot W5UDS0), frog *sst* (RefSeq XP_040205526.1), chicken *sst* (Uniprot P33094), mouse *Sst* (Uniprot P60041), human *SST* (Uniprot P61278). **(N)** Zebrafish *tac3a*-derived peptides detected in EECs. Alignment of the primary amino acid sequence of Tachykinin 3 includes zebrafish (Uniprot E9QE01), goldfish (Uniprot T2CZA2), catfish (Uniprot I4IY93), trout (Uniprot A0A8C7RBE8), salmon (Uniprot I4IY94), mouse (Uniprot P55099), rat (Uniprot A6HQX0), human (Uniprot Q9UHF0). **(O)** Zebrafish *vipb-*derived peptides detected in EECs. Peptides aligning to multiple isoforms of *vipb* were detected. Alignment of the primary amino acid sequence of Vasoactive intestinal peptides protein include zebrafish isoform 1 (Uniprot A0A8M9Q6Y9), zebrafish isoform 2 (Uniprot B0LF71), goldifsh (Uniprot A0A6P6NBR0), catfish (Uniprot A0A2D0RS13), salmon (Uniprot A0A1S3PEV), trout (Uniprot A0A060X5U6), chicken (Uniprot P48143), mouse (Uniprot A0A0R4J003), rat (Uniprot A0A090AX27), human (Uniprot P01282). **(P)** Zebrafish *ins*-derived peptides detected in EECs. Alignment of the primary amino acid sequences of Insulin protein include zebrafish (Uniprot B2GSI0), stickleback (Ensembl ENSGACT00000066960.1), tilapia (Uniprot I3IUZ1), herring (Ensembl ENSCHAT00020053570.1), xenopus (Uniprot F6QRS1), chicken (Uniprot P67970), mouse (Uniprot P01326), rat (Uniprot P01323), human (Uniprot P01308). **(Q)** Zebrafish *pcsk1nl*-derived peptides detected in EECs. **(R)** Zebrafish *pcsk1*-derived peptides detected in EECs. **(S)** Zebrafish *pcsk2*-derived peptides detected in EECs. **(T)** Zebrafish *scg2a*-derived peptides detected in EECs. **(U)** Zebrafish *scg2b*-derived peptides detected in EECs. **(V)** Zebrafish *scg3-*derived peptides detected in EECs. **(W)** Zebrafish *scg5*-derived peptides detected in EECs. **(X)** Zebrafish *scgn*-derived peptides detected in EECs.

**S4 Fig. Example raw peptidomic data of acylated ghrelin and manually curated peptides.**

**(A)** Fragmentation spectrum identifying octanolylated ghrelin peptide. **(B)** Fragmentation spectrum identifying decanolylated ghrelin peptide. **(C)** Chromatogram data showing overlapping peaks for double, triple, and quadruple charged YGGFLRKFGPK peptide manually identified from the Pdyn protein. **(D)** Chromatogram data showing overlapping peaks for isotopologues M, M+1, and M+2 of the +3 charged peptide. Peak areas are quantified below and mass error for each is shown. **(E)** Expression in the joint larval and adult intestinal dataset of hormone genes commonly enriched in the pancreatic islet.

**S5 Fig. Creation of *ghrl:QF2* and *ngn3:QF2* lines.**

**(A)** Schematic representation of cloning approach to generating *ngn3:QF2* and *ghrl:*QF2 reporters. **(B)** mVISTA [153,154] alignment of the *ghrl* locus in six closely related *Danio* species. *Danio rerio* *ghrl* annotation is shown at the top and is used as the reference. Dashed lines mark the highly conserved 500 base pair region upstream of the *ghrl* transcriptional start site that was cloned for the *ghrl:QF2* reporter. **(C)** Staining with anti-Ghrelin antibody and co-labeling with *ghrl:QF2* reporter in the **(D)** intestine and **(E)** pancreatic islet. **(F)** Staining with anti-Ghrelin antibody and co-labeling with *ghrl:QF2* reporter in the **(G)** intestine in another sample. **(H)** Quantification of *ghrl*+ cells overlap with the pan-EEC reporter *neurod1:RFP* across larval development. Each dot represents an individual fish. **(I)** A representative image of a 5dpf *ghrl:QF2* fish with a highlighted example of a *ghrl*+; *neurod1-* cell. **(J)** Schematic of where representative cross sections were taken to evaluate the *ghrl:QF2* reporter in adult intestines **(K)** Representative image of a proximal intestinal section showing *ghrl*+; *neurod1*+ cells with classical EEC morphology. **(L)** Representative image of *ghrl*+; *neurod1+* cell with classical EEC morphology in distal intestinal section. **(M)** Quantification of *ngn3*:*QF2* reporter overlap with *neurod1:RFP* reporter across larval development. Each dot represents an individual fish. **(N)** A representative image of a 6dpf *ngn3:QF2* reporter fish with a highlighted example of a *ngn3*+; *neurod1-* cell. All scale bars are 100 μm.

**S6 Fig. EEC subtype reporter imaging across larval development.**

Representative images showing subtype distribution and overlap with pan-EEC *neurod1* reporter at 3, 4, and 5dpf for **(A)** *ngn3* reporter, **(B)** *ghrl* reporter, **(C)** anti-PYY antibody, **(D** *gcga* reporter, **(E)** anti-CCK antibody, **(F)** *trpa1b* reporter. Scale bars are 100 μm.

**S7 Fig. EEC subtype reporter cell counts across larval development.**

The percent of *neurod1*+ cells that label with the subtype reporter and the raw counts of subtype numbers per each equal-length quarter of the gut is shown for 3, 4, 5, and 6 days post fertilization (dpf) fish in the **(A)** *ngn3* reporter, **(B)** *ghrl* reporter, **(C)** anti-PYY antibody **(D)** *gcga* reporter **(E)** anti-CCK antibody, and **(F)** *trpa1b* reporter. Each dot represents a fish.

**S8 Fig. *ngn3:Cre* induced ablation leads to similar loss of *neurod1*+ cells as *ngn3:QF2; QUAS:Cre.***

**(A)** Schematic of *ngn3:Cre* induced *ngn3+* cell ablation. **(B)** Total counts and regional counts of *neurod1*+ cells in the *ngn3:Cre* ablated and control fish. Each dot represents a 6 day post fertilization fish. Statistical significance was calculated by unpaired *t* test for total cell numbers and by two-way ANOVA for regional analysis. Significance annotations are as follows: ns (p>0.05), * (p<0.05), ** (p<0.01), *** (p<0.001), **** (p<0.0001).

**S9 Fig. Loss of *ghrl* does not impact *neurod1+* EEC numbers.**

**(A)** Schematic of the *ghrl* locus with arrow heads marking the sites targeted with CRISPR guide RNAs. **(B)** Genotyping and control gels run with 1 kb+ NEB ladder and 3 mutant and 3 wildtype samples. Amplification with genotyping primers (red arrows) across the 804 base pair region results in a 300 base pair band due to deletion in mutants (lanes 1-3). Wildtype samples (lanes 4-6) do not amplify due to the highly repetitive nature of intron 2, but all samples amplified with control primers amplifying at a non-affected locus (*mttp* gene). **(C)** Total counts and regional counts of *neurod1+* cells in wildtype (heterozygous) and *ghrl* mutant (homozygous) fish. Each dot represents a 6 day post fertilization fish. Statistical significance was calculated by unpaired *t* test for total cell numbers and by two-way ANOVA for regional analysis.

**S10 Fig. Raw images of gels used in publication.**
